# Exposure to sounds during sleep impairs hippocampal sharp wave ripples and memory consolidation

**DOI:** 10.1101/2023.11.22.568283

**Authors:** Karla Salgado-Puga, Gideon Rothschild

## Abstract

Sleep is critical for the consolidation of recent experiences into long-term memories. As a key underlying neuronal mechanism, hippocampal sharp-wave ripples (SWRs) occurring during sleep define periods of hippocampal reactivation of recent experiences and have been causally linked with memory consolidation. Hippocampal SWR-dependent memory consolidation during sleep is often referred to as occurring during an “offline” state, dedicated to processing internally generated neural activity patterns rather than external stimuli. However, the brain is not fully disconnected from the environment during sleep. In particular, sounds heard during sleep are processed by a highly active auditory system which projects to brain regions in the medial temporal lobe, reflecting an anatomical pathway for sound modulation of hippocampal activity. While neural processing of salient sounds during sleep, such as those of a predator or an offspring, is evolutionarily adaptive, whether ongoing processing of environmental sounds during sleep interferes with SWR-dependent memory consolidation remains unknown. To address this question, we used a closed-loop system to deliver non-waking sound stimuli during or following SWRs in sleeping rats. We found that exposure to sounds during sleep suppressed the ripple power and reduced the rate of SWRs. Furthermore, sounds delivered during SWRs (On-SWR) suppressed ripple power significantly more than sounds delivered 2 seconds after SWRs (Off-SWR). Next, we tested the influence of sound presentation during sleep on memory consolidation. To this end, SWR-triggered sounds were applied during sleep sessions following learning of a conditioned place preference paradigm, in which rats learned a place-reward association. We found that On-SWR sound pairing during post-learning sleep induced a complete abolishment of memory retention 24 h following learning, while leaving memory retention immediately following sleep intact. In contrast, Off-SWR pairing weakened memory 24 h following learning as well as immediately following learning. Notably, On-SWR pairing induced a significantly larger impairment in memory 24 h after learning as compared to Off-SWR pairing. Together, these findings suggest that sounds heard during sleep suppress SWRs and memory consolidation, and that the magnitude of these effects are dependent on sound-SWR timing. These results suggest that exposure to environmental sounds during sleep may pose a risk for memory consolidation processes.

## INTRODUCTION

Sleep is critical for memory consolidation- the stabilization of recent labile memory traces into long-term memories. The consolidation of episodic, spatial, and contextual information into long-term memories is strongly dependent on the hippocampus (Diekelmann & Born, 2010; Frankland & Bontempi, 2005; O’Keefe & Nadel, 1978; Scoville & Milner, 1957). A key neurophysiological mechanism implicated in hippocampal-dependent memory consolidation are sharp-wave ripples (SWRs). SWRs are brief bursts of high-frequency oscillations that originate in the hippocampus and are prominent during awake quiescence and non-rapid eye movement (NREM) sleep (Buzsaki, 1989, 2015). SWRs reflect periods of hippocampal-cortical communication (Buzsaki, 2015; Diekelmann & Born, 2010; Ego-Stengel & Wilson, 2010; Geva-Sagiv et al., 2023; Geva-Sagiv & Nir, 2019; Girardeau, Benchenane, Wiener, Buzsaki, & Zugaro, 2009; Jadhav, Rothschild, Roumis, & Frank, 2016; Ji & Wilson, 2007; Peyrache, Khamassi, Benchenane, Wiener, & Battaglia, 2009; Rothschild, Eban, & Frank, 2017; Squire, Genzel, Wixted, & Morris, 2015; Varela, Kumar, Yang, & Wilson, 2014; Varela & Wilson, 2020b; Wierzynski, Lubenov, Gu, & Siapas, 2009), during which reactivation of recent awake experiences occurs (Diba & Buzsaki, 2007; Foster, 2017; Skaggs & McNaughton, 1996; Wilson & McNaughton, 1994). Moreover, blocking SWRs using electrical stimulation impairs learning and memory (Ego-Stengel & Wilson, 2010; Girardeau et al., 2009; Jadhav, Kemere, German, & Frank, 2012). Together, SWRs are believed to be a critical neurophysiological mechanism for the emergence of long-term memories (Brodt, Inostroza, Niethard, & Born, 2023).

Although the sleeping brain is generally considered to be in an “offline” state, dedicated to processing internally generated activity patterns such as SWRs, the brain is not fully disconnected from the environment in this state. In particular, sounds heard during sleep are processed by a fully functional and highly active auditory system (Edeline, Dutrieux, Manunta, & Hennevin, 2001; Hayat et al., 2022; Issa & Wang, 2008; Nir, Vyazovskiy, Cirelli, Banks, & Tononi, 2015; Pena, Perez-Perera, Bouvier, & Velluti, 1999; Sela, Krom, Bergman, Regev, & Nir, 2020). The ability of the auditory system and the circuits it communicates with to process sounds during sleep is evolutionarily adaptive, as it supports rapid awakening in response to sounds of approaching predators or offspring calls (Velluti, 1997). However, many sounds processed by the brain during sleep are not behaviorally relevant. In particular, it is estimated that more than 20% of people living in major urban environments are regularly exposed during nighttime to the sounds of household appliances, outdoor traffic or other neighborhood noise (Brink, Omlin, Muller, Pieren, & Basner, 2011; Chepesiuk, 2005; Europe, World Health Organization, Hurtley, & World Health Organization. Regional Office for, 2009; Fiedler & Zannin, 2015; Fritschi & World health Organization. Regional Office for, 2011; Jarosinska et al., 2018). Yet, whether ongoing processing of sounds during sleep interferes with sleep-dependent cognitive processes and may therefore come at a cost for the process of memory consolidation, remains largely unexplored.

Direct projections from the auditory system to major hippocampal input regions, namely the perirhinal cortex and the lateral entorhinal cortex (Budinger & Scheich, 2009; Furtak, Wei, Agster, & Burwell, 2007; Insausti, Herrero, & Witter, 1997; Kerr, Agster, Furtak, & Burwell, 2007; Mascagni, McDonald, & Coleman, 1993; Steward, 1976), form an anatomical pathway by which incoming sounds during sleep can modify hippocampal activity. Indeed, functional studies have demonstrated that sounds can influence hippocampal activity (Billig, Lad, Sedley, & Griffiths, 2022; Kraus & Canlon, 2012). For example, during wakefulness, hippocampal place cells have been shown to encode unpredictable sounds, behaviorally relevant sounds or those representing a continuous acoustic space (Aronov, Nevers, & Tank, 2017; Itskov, Vinnik, Honey, Schnupp, & Diamond, 2012; Moxon et al., 1999; Sakurai, 2002; Xiao, Liu, Xu, Gan, & Xiao, 2018). During sleep, presenting sounds previously associated with a spatial location can bias hippocampal replay towards that location (Bendor & Wilson, 2012). In humans, presenting sounds during sleep that had previously been associated with other stimuli can strengthen memory retention of those stimuli upon awakening (Antony, Gobel, O’Hare, Reber, & Paller, 2012; Nicola Cellini & Alessandra Capuozzo, 2018; Creery, Oudiette, Antony, & Paller, 2015; Lewis & Bendor, 2019; Oudiette & Paller, 2013; Rudoy, Voss, Westerberg, & Paller, 2009). However, whether the ongoing processing of sounds that are unassociated with existing memories during sleep interferes with hippocampal SWR-dependent memory consolidation remains unknown.

To address this gap, we investigated the neural and behavioral influences of sound exposure during sleep in rats. To this end, we presented sounds to rats during sleep after spontaneous behavior and after learning paradigms and recorded the influence of sounds on hippocampal activity and memory consolidation. Furthermore, using a closed-loop system to present sounds during or following SWRs, we tested whether sound-induced modulation of neural activity and memory is limited to sounds occurring during SWRs. We find that exposure to sounds during sleep impairs SWRs and behaviorally measured memory consolidation. These effects were stronger, but were not limited to, sounds occurring during SWRs. Our results suggest that exposure to environmental sounds during sleep may pose a risk to memory functions.

## RESULTS

### Sounds during sleep suppress SWRs in a timing-dependent manner

To determine the consequence of exposure to sounds during sleep on hippocampal SWRs, we first presented brief (50 ms) low-intensity broad-band noise (BBN) stimuli during or following SWRs in naturally sleeping rats. To this end, SWRs were recorded from the dorsal CA1 region of the hippocampus during a 2 h sleep session following spontaneous behavior, and a closed-loop system was used to detect SWRs in real-time, allowing presentation of sounds during or following SWRs (**Figure 1A**). During the first hour of sleep, SWRs were detected, but no sound stimulus was given (No Stimulation, “NS”). During the second hour, a BBN sound stimulus was triggered by every detected SWR in sessions with either of two protocols: immediately following SWR onset detection, referred to as “On-SWR” (**Figure 1B**), or 2 seconds after SWR detection, referred to as “Off-SWR” (**Figure 1C**).

**Figure 1.**
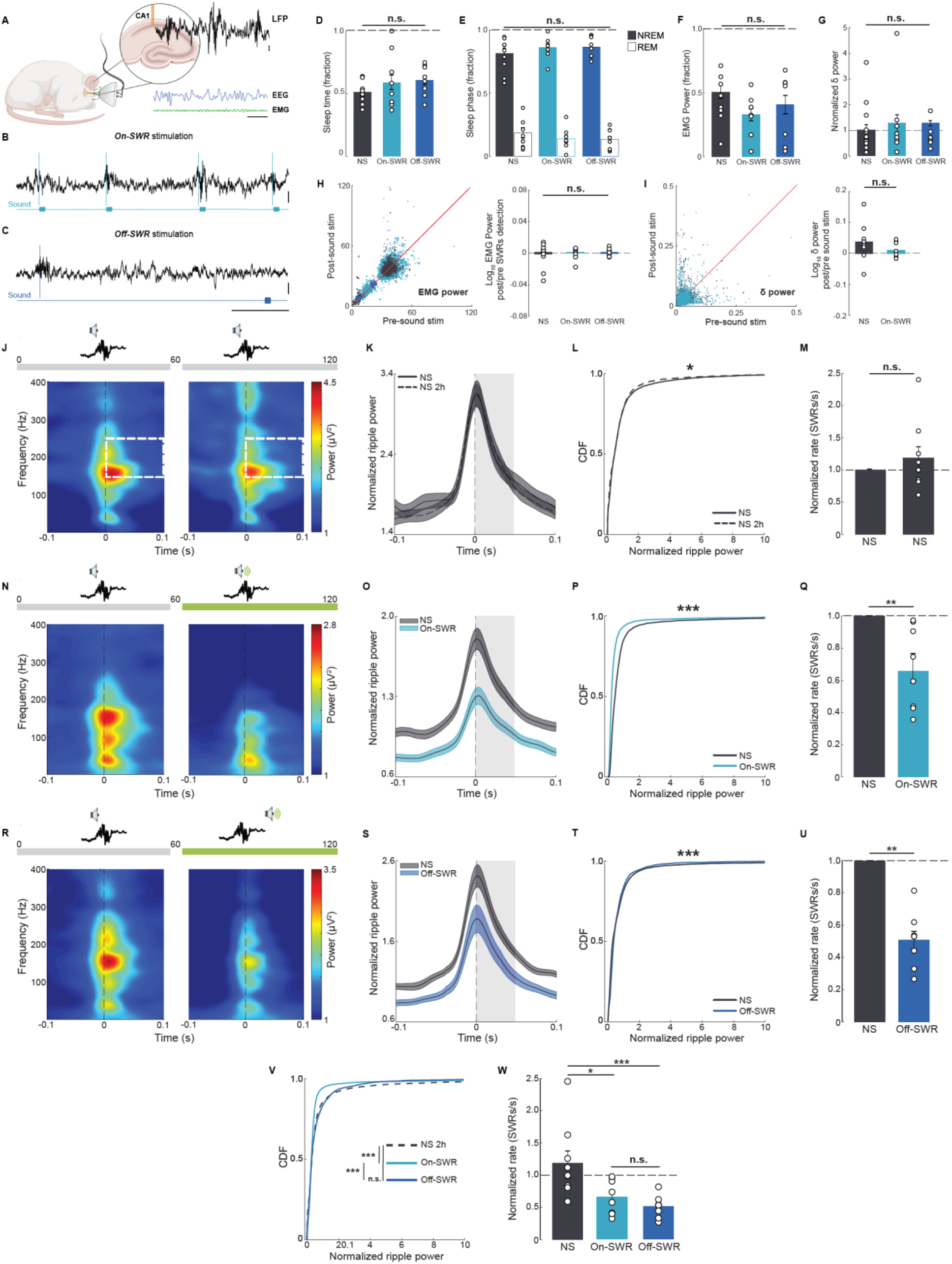
Exposure to non-waking sounds during sleep reduces ripple power and rate in a timing-dependent manner. **A-C.** The closed loop system for SWR-triggered sound stimulation. **A**. Illustration of hippocampal LFP recordings combined with EEG and EMG recordings in naturally sleeping rats. **B**. Sound stimulation paired to SWR onset (On-SWR). **C**. Sound stimulation delayed 2 seconds after SWR detection (Off-SWR). **D-I.** Sleep attributes of groups that had no sound stimulation (NS), sound paired to SWRs (On-SWR), or sound delayed 2s after SWR detection (Off-SWR). There were no significant differences among the groups in amount of sleep time (**D**, KW test p = 0.4394), fraction of time per sleep phase (**E**, KW test, p = 0.8573), EMG power per session (**F**, KW test, p = 0.1283), and normalized Delta power during NREM per session (**G**, NS group mean power = 1.0; KW test, p = 0.8564**). H-I**. PeriSound EMG and Delta power analysis. **H**, The EMG power 5 seconds before and after sound stimulation did not show significant difference between groups (KW test, p = 0.8421). **I**, The Delta power 5 seconds after sound stimulation from NS and On-SWR groups were not significantly different from the 5 seconds before sound stimulation (sample WSR test, p = 0.1484 and p = 0.5469, respectively). Neither there was a significant difference between NS and On-Rip groups (MW-U test, p = 0.0727). **J-X**. SWR recordings for 2 hours of sleep after an hour of spontaneous behavior. During the first hour SWRs were detected but no sounds were given (NS condition). During the second hour either no sound (NS), On-SWR, or Off-SWR sound stimulation was given. **J**, SWR spectrograms from the first and the second sleep hour with no sound stimulation (NS). The white dashed rectangles denote the time-frequency region of the spectrogram used for ripple power quantification. **K**, Normalized ripple power representation shown in the spectrograms. The gray area indicates the time frame of the sound stimulation if given. **L**, Cumulative distribution of the normalized ripple power quantified from the area represented in the white rectangle shown on the spectrograms. Ripple power is slightly but significantly reduced (0.9393 ± 0.034) compared to ripple power during the first hour of sleep (NS, mean power = 1.0, WSR test, *, p = 0.0007). **M**, Normalized SWR rates did not differ across the two hours (1.1887 ± 0.1868 from NS control condition (mean rate = 1.0), WSR test, p = 0.7344). **N-Q**, Same representation as in **J-M** but for the group with On-SWR sound stimulation during the second hour of sleep. **R-U**, Same representation as in **J-M** but for the group with Off-SWR sound stimulation during the second hour of sleep. Notice that both On-SWR and Off-SWR stimulations significantly reduced the ripple power (0.6861 ± 0.0394, (WSR test, *, p = 1.0786e^-118^, (**P**)) and 0.7116 ± 0.0198, (WSR test, *, p = 1.3006e^-105^, (**T**)), respectively) and rate (0.6563 ± 0.0953 (WSR test, *, p = 0.0078, (**Q**)) and 0.5082 ± 0.0538 (WSR test, *, p = 0.0078, (**U**)), respectively) compared to the NS condition. **V**, On-SWR stimulation induced a significant reduction in ripple power compared to the second hour of the NS group and the Off-SWR group (KW test, p = 9.4942e^-09^; T-K test, *, p = 1.5491e^-07^ and *, p = 2.7534e^-06^, respectively). Note that, despite the Off-SWR stimulation reducing ripple power, this effect was not significantly different from the second hour of the NS group (T-K test, p = 0.3061). **W**, On-SWR and Off-SWR stimulations significantly reduce ripple rate compared to the NS condition (KW test, p = 0.0025; T-K test, *, p = 0.0111 and, *, p = 0.0006, respectively). There was no significant difference between On-SWR and Off-SWR stimulations (T-K test, p = 0.3282). Data are presented as mean ± SEM, (n = 8-9 rats/group).

We used a BBN intensity of 50 dB, which is well above the rats’ hearing threshold (Borg, 1982) but which pilot experiments identified as not inducing wakefulness. We confirmed this by comparing key sleep attributes in the NS sessions to those of the On-SWR and Off-SWR sessions. The fraction of time spent asleep (**Figure 1D**), the fraction of sleep time spent in NREM and REM (**Figure 1E**), EMG power during sleep (**Figure 1F**) and EEG Delta power during sleep (**Figure 1G**) did not differ between the NS, On-SWR and Off-SWR sessions. Moreover, comparison of the EMG power and the Delta power 5s before and after sound stimulation did not show significant differences between groups (**Figure 1H-I**).

Next, we examined the influence of sound exposure on SWRs by comparing SWR statistics in the first hour of sleep (always NS) and the second hour of sleep (NS, On-SWR or Off-SWR). We normalized the power of each ripple to the mean ripple power in the first hour of sleep. Ripple power of SWRs showed a small (6.07%) but significant reduction between the first and second hour in the NS condition (**Figure 1J-L**), consistent with previous studies (Eschenko, Ramadan, Molle, Born, & Sara, 2008), while SWR rates did not significantly differ (**Figure 1M**). In contrast, pairing BBN to SWRs in the On-SWR condition caused a large (31.39%) and significant reduction in ripple power (**Figure 1N-P**), as well as a reduction in SWR rate (**Figure 1Q**). Interestingly, aligning ripple power to sound onset revealed that some of the reduction in ripple power was evident before sound onset, i.e., after the preceding sound (**Figure 1O**), reflecting a lasting effect of sounds on ripple power. Consistent with this finding, BBN presented 2 s following SWR onset in the Off-SWR condition also caused a significant reduction in ripple power (28.84%, **Figure 1R-T**) and SWR rate (**Figure 1U**), suggesting that the influence on SWRs persists beyond the immediate sound period.

To determine whether sounds occurring during SWRs have a stronger influence on SWR power and rate than those occurring between SWRs, we compared the reduction in ripple power from the first to the second hour of sleep across conditions (**Figure 1V, W**). We found that the On-SWR paradigm caused a significantly larger reduction in ripple power as compared to that of the Off-SWR paradigm (**Figure 1V**). The influence of On-SWR and Off-SWR paradigms on ripple rate did not significantly differ (**Figure 1W**). Other SWR features such as the ripple’s peak frequency or duration were not significantly modified by either of the sound stimulation protocols (**Supplementary Figure 1A, B**). Together, these findings demonstrate that mild, non-waking sounds heard during sleep suppress SWRs when sounds occur during or following SWRs, and that this suppression is stronger when it occurs during SWRs.

**Supplemental Figure 1.**
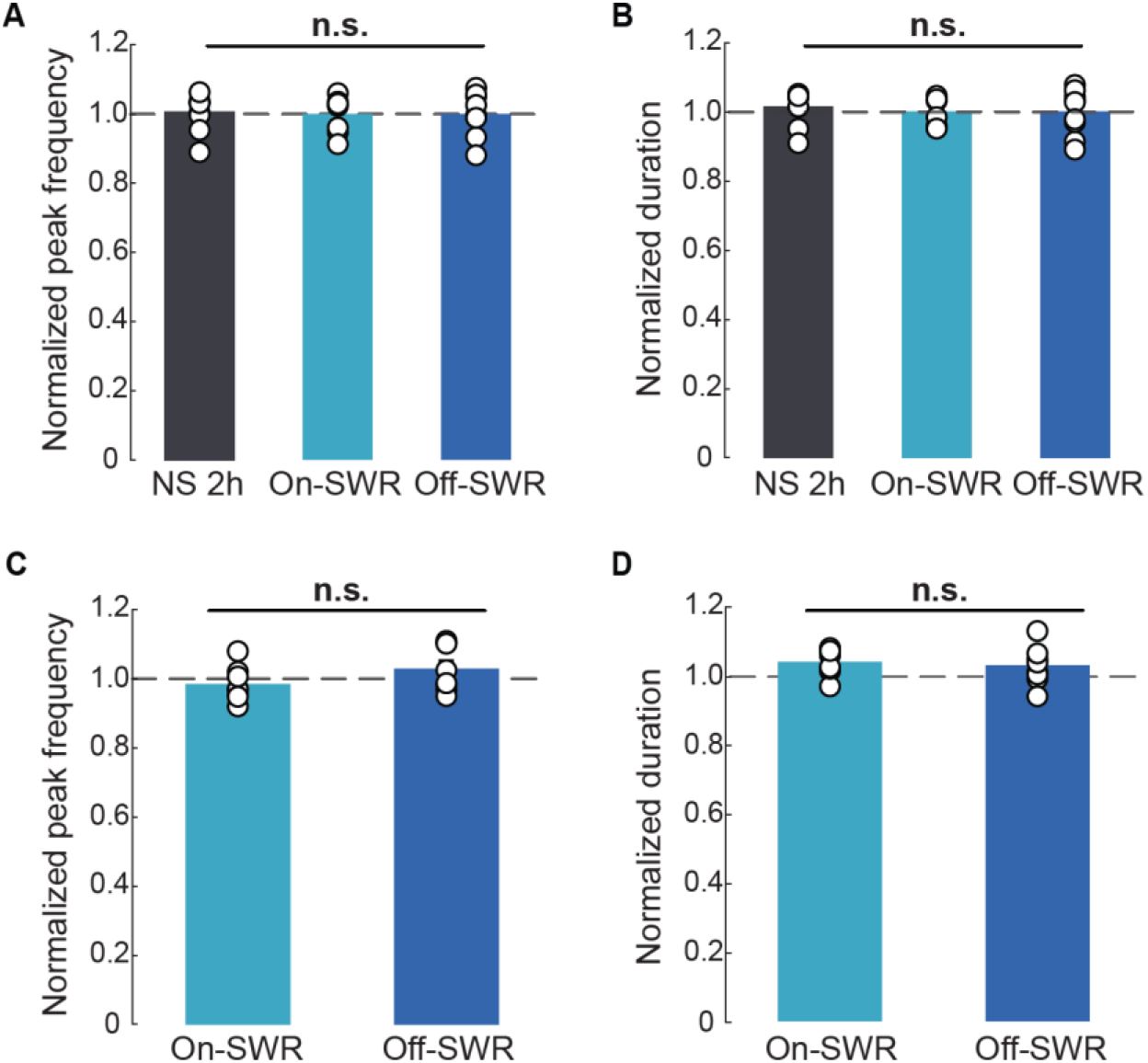
On-SWR and Off-SWR sound stimulation do not change SWRs peak frequency and duration. **A**. Normalized ripple peak frequency means comparison of the second hour of sleep with no sound (NS 2h, 1.005 ± 0.0177 from NS control condition (mean frequency = 1.0)), On-SWR (0.9979 ± 0.0174) or Off-SWR (1.0223 ± 0.0212) sound stimulation. Note that there is no significant difference between the first sleep hour (NS condition) and the second hour with no stimulation (WSR test, p = 0.9102), On-SWR (WSR test, p = 0.9453), or Off-SWR (WSR test, p = 0.1641) sound stimulation. Neither is there a difference between groups (KW test, p = 0.4508). **B**, Normalized SWR duration means comparison of the second hour of sleep on NS (01.014 ± 0.0162), On-SWR (1.0072 ± 0.0139), and Off-SWR (0.9682 ± 0.0211) conditions did not show significant difference compared to the first hour of sleep (WSR test, p = 0.5469, p = 0.5460, and p = 0.6523, respectively). Comparison between groups also showed no significant difference (KW test, p = 0.4024). **C.** Comparison of the normalized peak frequency of both On-SWR (0.9787 ± 0.0176) and Off-SWR (1.0230 ± 0.0235) sound stimulation did not show a significant difference compared to the NS group (KW test, p = 0.5298). **D.** Comparison of the normalized duration of On-SWR (1.039 ± 0.0138) and Off-SWR (1.0280 ± 0.0198) sound stimulation did not show a significant difference compared to the NS group (KW test, p = 0.1575). Data are presented as mean ± SEM, (n = 8-10 rats/group).

### Exposure to sounds during SWRs in sleep impairs memory

As SWRs during sleep are known to support memory consolidation (Ego-Stengel & Wilson, 2010; Fernandez-Ruiz et al., 2019; Girardeau et al., 2009; van de Ven, Trouche, McNamara, Allen, & Dupret, 2016), we next tested whether the sound-induced suppression in SWRs we observed is also associated with impaired memory. To this end, we used a hippocampal-dependent appetitive conditioned place preference (CPP) paradigm (Roy et al., 2017; Trouche et al., 2019) (**Figure 2A**). In the first stage of this task, performed in a setup with a center home chamber and two external chambers, the rats’ innate preference between each of the external chambers was recorded based on the amount of time spent in each. In the learning session of this task, rats learned to associate their less-preferred chamber with sucrose solution reward while water was placed in the other external chamber. The learning session was followed by a ∼3 h sleep session in the middle chamber, during which varying acoustic protocols were presented (see below). Memory retention was subsequently evaluated as the relative amount of time spent in the rewarded chamber. Memory was evaluated at two time points: immediately following the 3 h sleep session when memory retention relies largely on short-term processes (I. Izquierdo et al., 1998; Lee, Everitt, & Thomas, 2004; Rudy & Sutherland, 2008; Schafe, Nadel, Sullivan, Harris, & LeDoux, 1999a; Tse et al., 2007), and at 24 h post learning, when memory relies on hippocampal-dependent memory consolidation (I. Izquierdo et al., 1998; Lee et al., 2004; Rudy & Sutherland, 2008; Schafe et al., 1999a; Tse et al., 2007). In the first experimental group, the On-SWR BBN pairing paradigm was applied during the post-learning sleep session (**Figure 2B**). Consistent with the data following spontaneous behavior, SWRs in the On-SWR condition had significantly weaker ripple power as compared to the NS condition (**Figure 2C-E**), while SWR rates were not significantly different (**Figure 2F**). Behaviorally, under baseline conditions in which no sounds were presented in the post-learning sleep session, animals displayed significant memory retention at both time points, as evidenced by a significant learning-dependent shift in preference to the reward-associated chamber (**Figure 2G**). However, subjecting animals to the On-SWR sound paradigm during post-learning sleep caused a striking and significant memory impairment at the 24 h timepoint (**Figure 2H, J**), while memory retention at the 3 h timepoint was intact (**Figure 2H, I**). Moreover, at the 24 h timepoint CPP scores showed no difference from the pre-learning scores (**Figure 2H**), demonstrating a complete abolishment of memory. These findings suggest that sounds occurring during SWRs in sleep suppress ripple power and impair memory consolidation.

**Figure 2.**
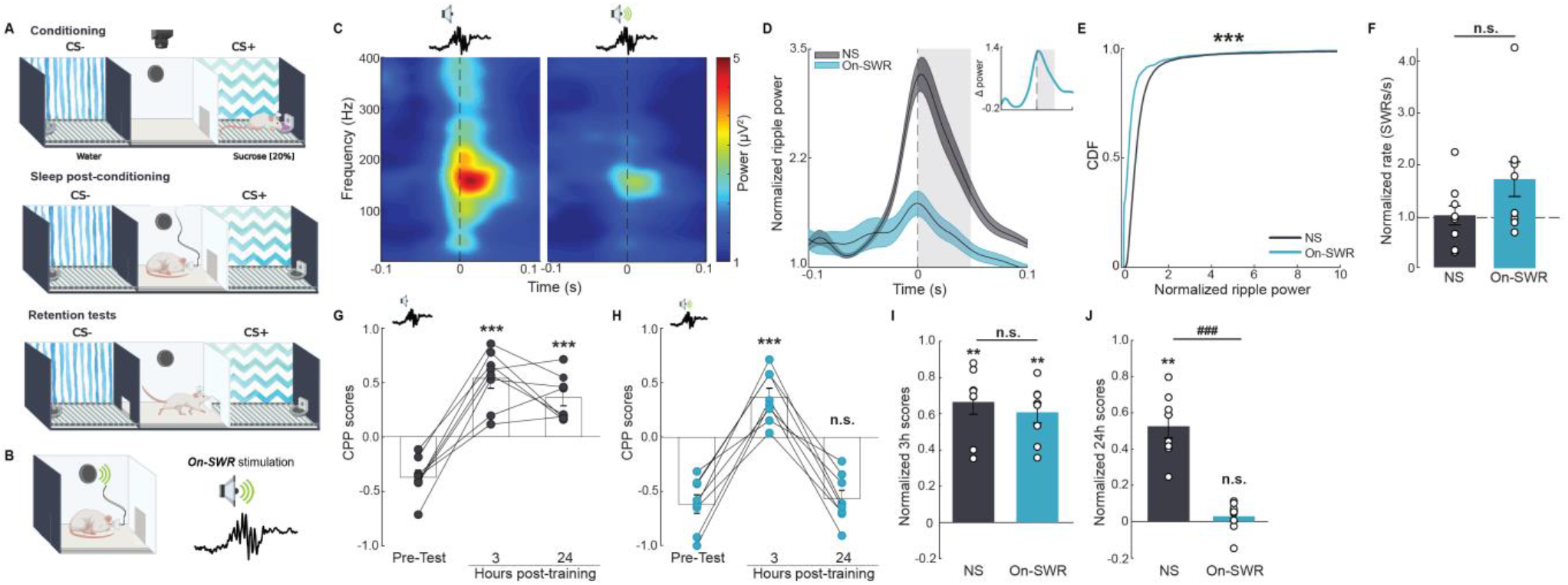
Sounds occurring during SWRs in sleep impair memory consolidation. **A.** Experimental design for the sound stimulation post-conditioning in the CPP task. **B.** The On-SWR sound stimulation protocol. **C.** Ripple spectrograms from groups NS and On-SWR. **D.** Normalized ripple power. **E.** Cumulative distribution of normalized ripple power. Note that On-SWR stimulation significantly reduced the ripple power (0.4962 ± 0.0130) compared to NS control group (mean power = 1.0, MW-U test, *, p = 0.0000). **F.** On-SWR stimulation did not significantly modify ripple rate compared to NS group (normalized mean rate 1.7218 ± 0.3362, MW-U test, p = 0.1949). **G-H.** CPP scores during Pre-Test, 3 h and 24 h post-training for the NS group (**G**) and the On-SWR group (**H**). CPP scores from both NS (0.5405 ± 0.0921) and On-SWR (0.3707 ± 0.0826) groups compared to the pre-Test scores (-0.3698 ± 0.0698 and -0.6175 ± 0.0852, respectively) showed a significant place preference shift 3 h post-conditioning (Friedman test, WSR post-hoc, *, p = 0.0078 and, *, p = 0.0078, respectively). In contrast, at 24 h post-conditioning, only the NS group showed a significant place preference shift (0.3623 ± 0.0698, WSR post-hoc, *, p = 0.0078, **G**), while the On-SWR group showed no significant difference to the pre-Test condition (-0.5674 ± 0.0789, WSR test, p = 0.2500, **H**). CPP scores at 24 h test also showed a significant reduction compared to the 3 h test (WSR test, *, p = 0.0078, **H**). **I-J**, Comparison of normalized scores for the 3 h (**I**) and 24 h (**J**) retention tests. Note that On-SWR stimulation induced a significant reduction of the normalized scores at the 24 h (0.0277 ± 0.0297, MW-U test, #, p = 1.5540e-^04^) but not at the 3 h (0.6058 ± 0.0559, MW-U test, p = 0.3823) retention test, compared to the NS group (0.5246 ± 0.0631 and 0.6649 ± 0.0675, respectively). For 3 h scores, sample WSR test, p = 0.0078 versus zero (zero considered as no memory). For 24 h scores, *, p = 0.0078 and p = 0.3125. Data are presented as mean ± SEM, (n = 8 rats/group).

### Sounds occurring outside SWRs in sleep weaken memory

Given that sound presentation in between SWRs caused a lingering reduction in the ripple power and rate of SWRs following spontaneous behavior (**Figure 1R-U**), we next tested whether it also induced memory impairments. We thus repeated the experiment described above but using the Off-SWR sound delivery paradigm in the post-learning sleep session (**Figure 3A**). As expected from the sleep data following spontaneous behavior (**Figure 1**), ripple power was significantly reduced as compared to the NS conditions (**Figure 3B-D**) and SWR rates were significantly lower (**Figure 3E**). Interestingly, we found that memory retention in the Off-SWR group was weaker compared to the NS group (**Figure 3F-I**). Consistent with the weakening of SWRs by Off-SWR sounds, these findings suggest that sounds occurring between SWRs also impair memory. Nevertheless, although weakened, a significant degree of memory retention was evident at both the 3 h and 24 h time points (**Figure 3G, I**). These results suggest that during NREM sleep, even sounds that occur largely outside of SWRs reduce SWR ripple power and decrease memory robustness.

**Figure 3.**
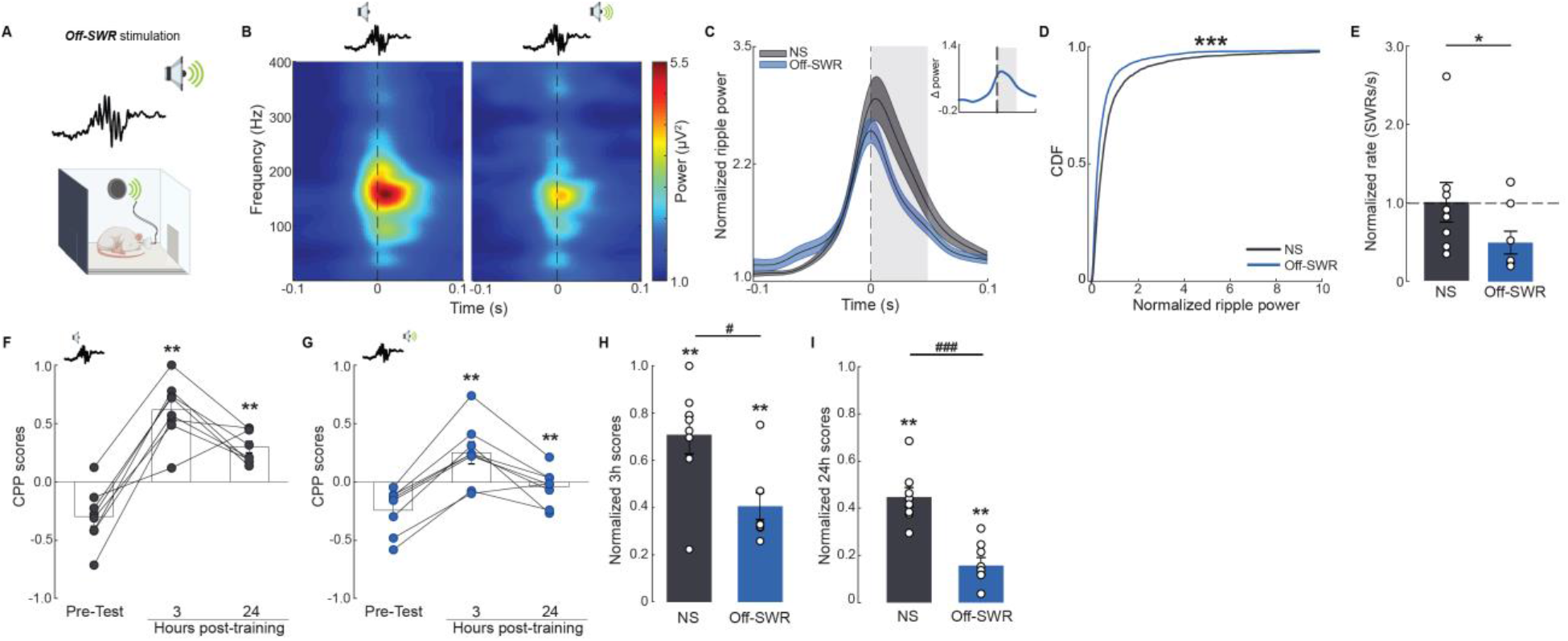
Sounds occurring during SWRs in sleep weaken memory consolidation. **A.** Sound stimulation protocol. **B.** SWR spectrograms from NS and Off-SWR groups. **C.** Normalized ripple power. **D.** Cumulative distribution of normalized ripple power. Off-SWR stimulation significantly reduced the ripple power (0.6387 ± 0.0204) compared to NS control group (mean power = 1.0, MW-U test, *, p = 4.1419e^-54^). **E.** Off-SWR stimulation significantly reduced SWR rate compared to NS group (normalized mean rate 0.4893 ± 0.1432, MW-U test, *, p = 0.0499). **F-G**, CPP scores during Pre-Test, 3 h and 24 h post-training for the NS (**F**) and Off-SWR (**G**) groups. CPP scores from both NS (0.6162 ± 0.0919) and Off-SWR (0.2485 ± 0.0942) groups compared to the pre-Test scores (-0.2976 ± 0.0858 and -0.2403 ± 0.0686, respectively) showed a significant place preference shift 3 h post-conditioning (Friedman test, WSR post-hoc, *, p = 0.0078 and, *, p = 0.0078, respectively). Similarly, at 24 h post-conditioning, both NS (0.2974 ± 0.0488) and Off-SWR (-0.0405 ± 0.0549) groups showed a significant place preference shift (WSR post-hoc, *, p = 0.0078 and, *, p = 0.0078, respectively). **H-I**, Normalized CPP scores at 3 h (**H**) and 24 h (**I**) retention tests, showing that Off-SWR stimulation significantly reduced both 3 h (0.4053 ± 0.0561, MW-U test, #, p = 0.0281) and 24 h (0.1557 ± 0.0355, MW-U test, #, p = 1.5540e-^04^) scores compared to the NS group (0.7071 ± 0.0802 and 0.4466 ± 0.0409, respectively), suggesting a weaker memory trace. For all scores, sample WSR test, *, p = 0.0078 versus zero (zero considered as no memory). Data are presented as mean ± SEM, (n = 8 rats/group).

The neural processes supporting memory consolidation can vary based on various task attributes, such as stimulus valence, stimulus novelty and task complexity (Pinto et al., 2019; Redondo et al., 2014; Schlüter, Hackländer, & Bermeitinger, 2019; Warden & Miller, 2010). Therefore, the susceptibility to memory disruption by sound exposure during sleep may also differ. Hence, to determine whether the sound-induced memory impairment we observed is task-specific, we carried out similar experiments using the Contextual Fear Conditioning (CFC) paradigm. CFC is a well-known hippocampal-dependent paradigm that differs from the CPP paradigm in both valence and complexity. Interestingly, we found that neither the On-SWR nor the Off-SWR sound protocols had a significant effect on memory consolidation of this task at either of the retention test timepoints (**Supplementary Figure 2C-F**). These findings demonstrate that the influence of sounds during sleep are task specific. Additionally, as CFC memory depends on intact post-training sleep (Graves, Heller, Pack, & Abel, 2003; Hagewoud et al., 2010) , these findings provide additional confidence that the impairment in memory retention of the CPP task did not stem from impaired sleep.

**Supplemental Figure 2.**
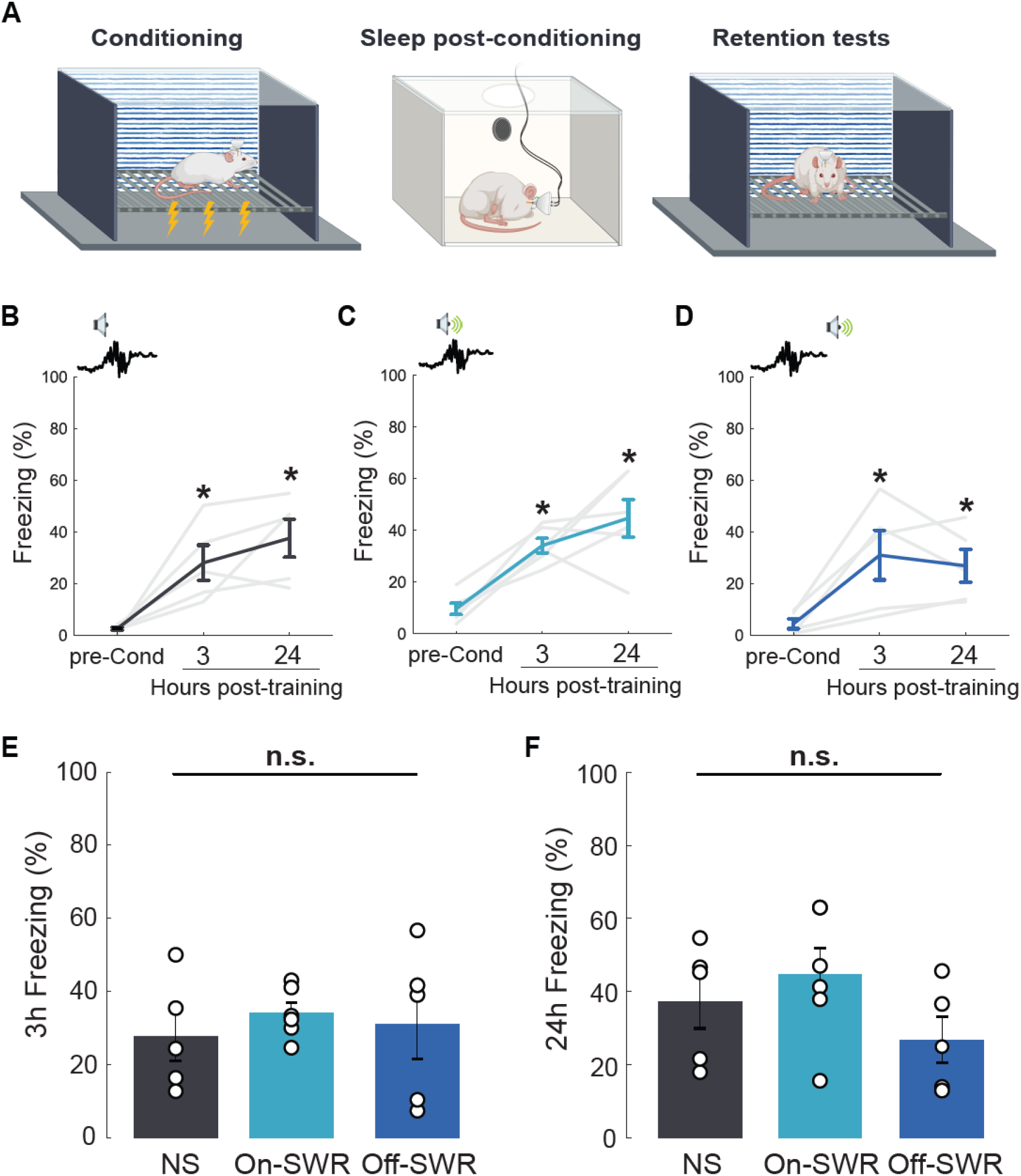
On-SWR and Off-SWR stimulation do not alter aversive memory consolidation. **A**, Experimental design for the sound stimulation post-conditioning in the CFC task. Sleep recordings post-training lasted 3-4h. Retention tests for the CFC task were done at 3 h and 24 h post-training. **B-D**, Proportion of freezing behavior during pre-Conditioning (pre-Cond), 3 h and 24 h retention tests for the groups NS (**B**), On-SWR (**C**), and Off-SWR (**D**). Note that all groups show a significant increase in freezing behavior at 3 h (27.75 ± 6.7947, 34.05 ± 2.8223, 31.0 ± 9.5493, respectively) and 24 h (37.27 ± 7.3169, 44.67 ± 7.2459, 26.87 ± 6.3671, respectively) post-conditioning compared to the pre-Conditioning (2.28 ± 0.6134, 9.60 ± 2.1610, 4.43 ± 1.9605, respectively; Friedman test, p < 0.05; WSR post-hoc test, *, p < 0.05). **E-F**, Comparison of the proportion of freezing behavior at 3 h (**E**) and 24 h (**F**) retention tests did not show a significant difference between groups (KW test, p = 0.7973 and p = 0.1806). Data are presented as mean ± SEM, (n = 5-6 rats/group).

### Sounds occurring during SWRs induce stronger neural and memory impairments than sounds occurring outside SWRs

Finally, we directly compared the neural and behavioral consequences of sound presentation in the On-SWR and Off-SWR paradigms during the post-learning sleep session. Consistent with our findings in sleep following spontaneous behavior, ripple power was significantly lower in the On-SWR paradigm than in the Off-SWR paradigm (**Figure. 4A, B**), suggesting that sounds that occur during SWRs have a more detrimental influence on ripple power. Interestingly, SWR rates were significantly lower in the Off-SWR paradigm as compared to the On-SWR paradigm (**Figure 4C**), suggesting that sounds have a differential influence on the power and rate of SWRs. As previously observed, the sound stimulation protocols did not modify the peak frequency or duration of SWRs (**Supplementary Figure 1C, D**). Finally, we compared the behavioral consequences of On-SWR and Off-SWR paradigms and found a differential effect in the different memory-retention time points. At the 3 h timepoint immediately after awakening, memory retention following the Off-SWR paradigm was significantly weaker than the On-SWR paradigm (**Figure 4D**), although animals in both the On-SWR and Off-SWR groups showed significant levels of memory retention (**Figure 2I** and **Figure 3H**). In contrast, at the 24 h timepoint, the On-SWR paradigm impaired memory formation significantly more than the Off-SWR paradigm (**Figure 4E**). These data are consistent with our finding that animals in the Off-SWR group showed significant memory retention at the 24 h timepoint (**Figure 3I**), while animals in the On-SWR group did not (**Figure 2J**). Together, these findings suggest that exposure to sounds during sleep has a detrimental influence on hippocampal SWRs and memory, and that the timing of sounds relative to SWRs has differential influences on the degree of these impairments.

**Figure 4.**
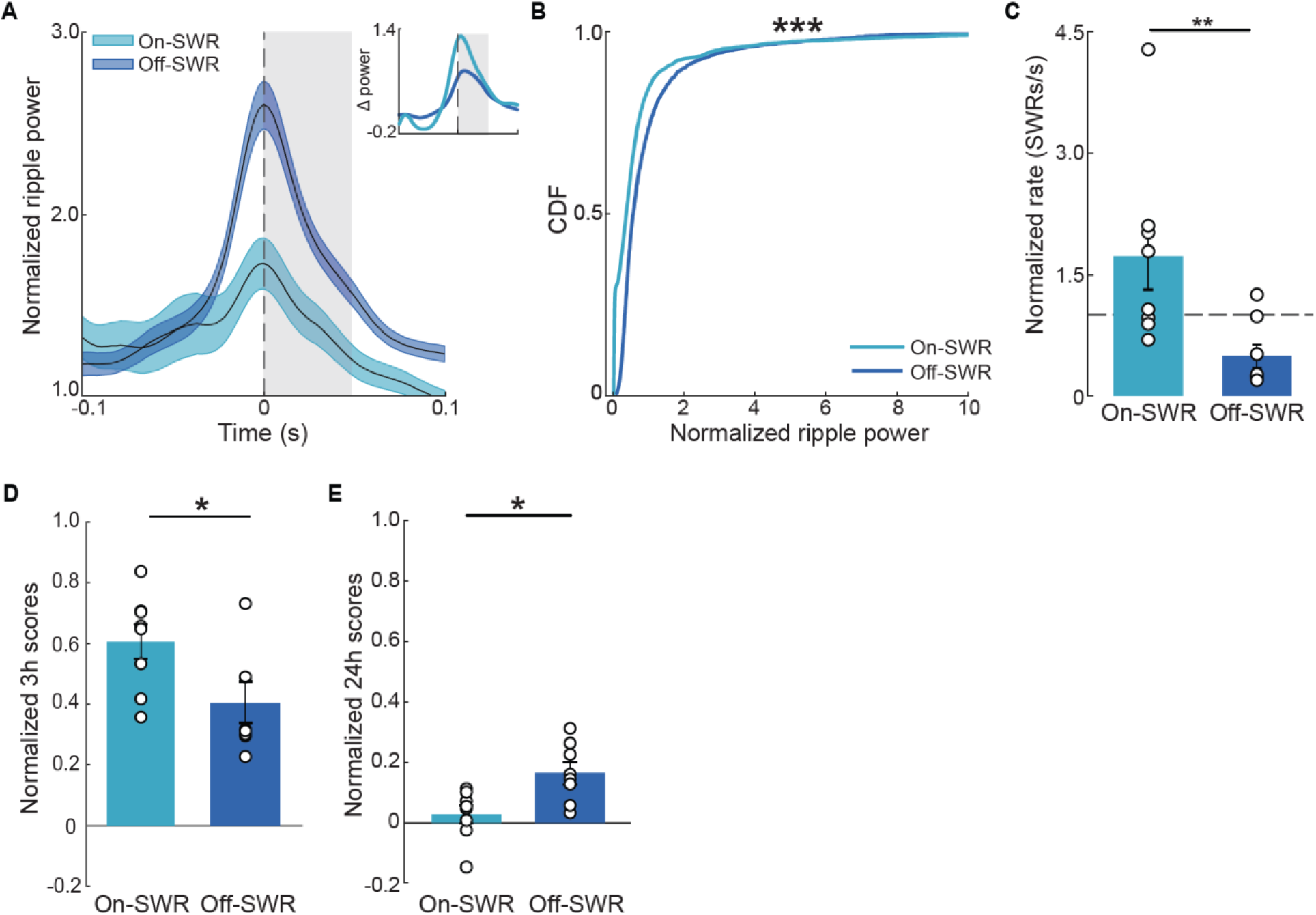
Sounds occurring during SWRs have greater effects on ripple power and memory than sounds occurring outside SWRs. **A.** Normalized ripple power of On-SWR (Figure 2C), and Off-SWR (Figure3B) groups. Inset, difference in normalized ripple power. **B.** Cumulative distribution of normalized ripple power. Note that ripple power suppression is significantly greater in the On-SWR group (0.4962 ± 0.0130) compared to the Off-SWR group (0.6387 ± 0.0204, MW-U test, *, p = 4.4146e^-136^). **C.** Off-SWR stimulation significantly reduced ripple rate (0.4893 ± 0.1432) compared to On-SWR stimulation (1.7218 ± 0.3362, MW-U test, *, p = 0.0070). **D-E.** Comparison of normalized scores for the 3 h (**D**) and 24 h (**E**) retention tests. Normalized CPP scores comparison at the 3 h retention test showed that On-SWR scores (0.6058 ± 0.0559) are significantly higher than Off-SWR (0.4053 ± 0.0561; MW-U test, *, p = 0.0028). In contrast, at the 24 h retention test On-SWR scores (0.0277 ± 0.0297) are significantly smaller compared to the Off-SWR scores (0.1557 ± 0.0355; MW-U test, *, p = 0.0207). Data are presented as mean ± SEM, (n = 8 rats/group).

## DISCUSSION

The auditory system continuously processes sounds during sleep (Edeline et al., 2001; Hayat et al., 2022; Issa & Wang, 2008; Nir et al., 2015; Pena et al., 1999; Sela et al., 2020) and communicates sound information to the medial temporal lobe via cortical and subcortical pathways (Budinger & Scheich, 2009; Furtak et al., 2007; Insausti et al., 1997; Kerr et al., 2007; Mascagni et al., 1993; Steward, 1976). However, whether processing of incoming sounds interferes with hippocampal-dependent memory consolidation has remained unknown. Our experiments reveal that exposure to sounds during sleep interferes with SWRs-a key biomarker of memory consolidation (Buzsaki, 2015; Diba & Buzsaki, 2007; Diekelmann & Born, 2010; Ego-Stengel & Wilson, 2010; Fernandez-Ruiz et al., 2019; Foster, 2017; Girardeau et al., 2009; Ji & Wilson, 2007; Peyrache et al., 2009; Rothschild et al., 2017; Skaggs & McNaughton, 1996; Squire et al., 2015; van de Ven et al., 2016; Wierzynski et al., 2009; Wilson & McNaughton, 1994), and with the consolidation of recent experiences into memory. Moreover, we found that sounds occurring during SWRs had the strongest influence on ripple power and memory retention 24 h after learning, but sounds occurring between SWRs also impaired these neural and behavioral measures. Together, our findings suggest that exposure to sounds during sleep impairs hippocampal-dependent memory consolidation in a task-specific manner.

SWRs define times of hippocampal replay (Diba & Buzsaki, 2007; Foster, 2017; Skaggs & McNaughton, 1996; Wilson & McNaughton, 1994) and hippocampal-cortical communication (Buzsaki, 2015; Diekelmann & Born, 2010; Ego-Stengel & Wilson, 2010; Geva-Sagiv et al., 2023; Geva-Sagiv & Nir, 2019; Girardeau et al., 2009; Ip et al., 2021; Ji & Wilson, 2007; Peyrache et al., 2009; Rothschild et al., 2017; Squire et al., 2015; Varela et al., 2014; Varela & Wilson, 2020b; Wierzynski et al., 2009), attributes which have identified SWRs as a candidate mechanism underlying the consolidation of recent experience into long-term memory representations. To test whether SWRs have a causal role in memory consolidation, previous studies used direct hippocampal commissure stimulation to interrupt SWRs following learning (Ego-Stengel & Wilson, 2010; Girardeau et al., 2009; Girardeau, Cei, & Zugaro, 2014; Michon, Sun, Kim, Ciliberti, & Kloosterman, 2019; Norimoto et al., 2018). These studies found that blocking SWRs using electrical stimulation impaired learning and memory formation. Our results show that SWR interruption and subsequent impairments in memory consolidation can also occur as a result of simple processing of sensory information during sleep. These findings suggest that “online” processing of incoming sounds during sleep may come at a cost for “offline” processing of internally generated activity patterns underlying memory consolidation.

We found that in sleep following spontaneous behavior, sounds occurring during SWRs (On-SWRs) as well as sounds occurring between SWRs (Off-SWRs) weakened ripple power, yet On-SWR sounds weakened it significantly more. Furthermore, both sound protocols caused a similar and significant reduction in SWR rates. These results and the known anatomical circuits point at likely underlying mechanisms. Sounds occurring during SWRs may weaken ripple power by generating activation throughout the auditory pathway which propagates via the entorhinal cortex or perirhinal cortex into the hippocampus (Budinger & Scheich, 2009; Insausti et al., 1997; Kerr et al., 2007; Steward, 1976), thereby interrupting with the synchronized activity in CA3 that is necessary for SWR generation (Memmesheimer, 2010; Schomburg et al., 2014; Ylinen et al.). However, weakening of ripple power by sounds occurring between SWRs (albeit to a lesser degree), as well as the reduction in SWR rate in both protocols, suggest that beyond their immediate effect, sounds weaken SWRs for hundreds of milliseconds or seconds after sound termination. Interestingly, a lingering effect of sounds on hippocampal activity during sleep has been previously shown (Bendor & Wilson, 2012). In this study, presentation of behaviorally relevant sounds during sleep influenced the content of hippocampal reactivation up to 10.8 seconds after sound presentation. In our data, the mechanism underlying this lingering effect may be explained within the abovementioned circuit, by the time period required for the recovery of synchronous network dynamics in CA3 following its interruption by sounds. Additionally, as the spiking patterns and oscillatory activity between SWRs can influence the content of activity during the SWRs (Chambers, Berge, & Vervaeke, 2022; Gridchyn, Schoenenberger, O’Neill, & Csicsvari, 2020; Hasselmo, 2003; Hasselmo & Stern, 2013), sounds interfering with pre-SWR activity may cause SWR weakening via this 2-stage process.

Another potential mechanism underlying the findings described above is cholinergic neuromodulation. Previous studies have found that cholinergic input from the medial septum (MS) into the CA3 field of the hippocampus suppresses SWRs and that this effect persists for at least hundreds of milliseconds following stimulation (Jarzebowski, Tang, Paulsen, & Hay, 2021; Vandecasteele et al., 2014). Furthermore, MS is activated by sounds and communicates sound information into the hippocampus (Miller & Freedman, 1993; Zhang et al., 2018). Thus, sounds heard during sleep may suppress SWRs via cholinergic input from the MS. Nevertheless, future studies are required to test these hypotheses and determine the detailed circuit mechanisms underlying sound-induced SWRs suppression.

During sleep following learning, hippocampal reactivation and hippocampal-cortical communication during SWRs are believed to be critical for the consolidation of recent experiences into long-term memories (Brodt et al., 2023; Buzsaki, 1989, 2015; Diekelmann & Born, 2010; Ego-Stengel & Wilson, 2010; Fernandez-Ruiz et al., 2019; Girardeau et al., 2009; Ji & Wilson, 2007; Peyrache et al., 2009; Rothschild et al., 2017; Squire et al., 2015; van de Ven et al., 2016; Wierzynski et al., 2009). We found that during post-learning sleep sessions, On-SWR and Off-SWR sounds induced a similar pattern of ripple power suppression as they did following spontaneous behavior. Interestingly, however, we found distinct effects of On-SWR and Off-SWR protocols on SWR rates following learning. While Off-SWR sounds induced a reduction in SWR rates as they did following spontaneous behavior, On-SWR sounds did not. Thus, there was a selective preservation of SWR rates in the On-SWR conditions post-learning. These findings are consistent with a previous study showing that following learning, but not following random behavior, compensatory mechanisms of SWR rates are engaged in response to SWR disruption (Girardeau et al., 2014). Nevertheless, the sound-induced reduction in ripple power following learning could interfere with memory consolidation and impair memory performance via the mechanisms described above. In particular, the reduction in ripple power following both the On-SWR and Off-SWR sounds suggests a reduced recruitment of neuronal firing in the CA1 output region of the hippocampus. This could reflect a reduction in the integrity of replay events that take place during the SWRs, and/or a reduction in the robustness of information flow from the hippocampus to cortical regions and the synaptic plasticity that this communication induces (Sadowski, Jones, & Mellor, 2016; Varela & Wilson, 2020a).

Memory is traditionally classified into short-term memory (STM), which lasts from seconds to a few hours, and long-term memory (LTM), which lasts from hours to days or longer (L. A. Izquierdo, Vianna, et al., 2000; Pignatelli et al., 2019; Roy et al., 2017). Several studies have established that STM and LTM rely on different neural mechanisms (I. Izquierdo et al., 1998; L. A. Izquierdo, Barros, et al., 2000; Lee et al., 2004; Poo et al., 2016; Rudy & Sutherland, 2008; Schafe et al., 1999a; Tse et al., 2007). For example, inhibiting the expression of Brain-Derived Neurotrophic Factor in the dorsal hippocampus impairs LTM, measured at 24 h post-training, while having no effect on STM, measured at 3 h post-training (Lee et al., 2004). Similarly, hippocampal protein synthesis inhibition before learning of a CFC paradigm caused LTM impairment 24 h post-conditioning while STM at the 1h post-conditioning was intact (Schafe, Nadel, Sullivan, Harris, & LeDoux, 1999b). Consistently, infusion of several protein kinase inhibitors into the hippocampus immediately after training in an inhibitory avoidance task had no effect over STM (tested 2 h post-training) but impaired LTM when tested 24 h post-training (Vianna et al., 2000; Walz et al., 2000). Accordingly, our findings reveal a differential effect of the On-SWR stimulation protocol on these stages of memory, wherein memory retention at the 3 h time point was intact (**Figure 2I**), while memory retention at 24 h was abolished (**Figure 2J**). These findings suggest that the On-SWR sound stimulation did not affect STM-related mechanisms but only those related to LTM formation. On the other hand, the deleterious effect of Off-SWR sound stimulation on memory retention at the 3 h timepoint (immediately following post-learning sleep) suggests that sounds outside SWRs may affect a different set of memory-related mechanisms. As previously stated, Off-SWR stimulation might be interfering with oscillatory events that happen outside of SWRs, which may be particularly important for STM.

The deleterious effects of sound on hippocampal function and memory are likely strongly dependent on the nature of the presented sound and its associated meaning. In the current study we focused on unfamiliar sounds associated with no behavioral meaning, as a model of the influence of environmental noise exposure. It is of interest to compare the current findings with those of a growing number of studies in recent years, which have examined the influence of presentation of sounds associated with a recently formed memory during sleep in rodents (Kronenberg, Milinski, Kruschke, & Hoz, 2022; Purple, Sakurai, & Sakaguchi, 2017) and in humans (Batterink, Creery, & Paller, 2016; Leminen et al., 2017). For example, presenting sounds with varying degrees of behavioral relevance during NREM sleep differentially affected sleep-associated oscillations in rodents (Kronenberg et al., 2022). In humans, targeted memory reactivation -TMR-studies describe strengthening of specific memories previously associated with the re-presented sound (Antony et al., 2012; Bar et al., 2020; N. Cellini & A. Capuozzo, 2018; Creery et al., 2022; Creery et al., 2015; Oudiette & Paller, 2013; Rudoy et al., 2009; Schechtman et al., 2021). Thus, it is likely that robust encoding of sound meaning (or lack thereof) during sleep strongly shapes the nature of the influence on memory consolidation.

Our findings of the deleterious effects of sound exposure during sleep on memory consolidation have potential broad public health relevance. Exposure to environmental sounds during sleep is highly prevalent in urban environments (Brink et al., 2011; Chepesiuk, 2005; Fiedler & Zannin, 2015; Hahad et al., 2022; Paul, Haan, Mayeda, & Ritz, 2019), but their influence on neural function and cognitive abilities is poorly understood. Epidemiological studies have found that long-term exposure to nighttime noise is associated with an increased risk for memory impairments, cognitive decline and dementia (Europe et al., 2009; Fritschi & World health Organization. Regional Office for, 2011; Jarosinska et al., 2018; Tzivian et al., 2016; Tzivian et al., 2017; Weuve et al., 2020). Whether repeated interruption of hippocampal activity as we observed in an acute exposure setting, eventually leads to lasting or permanent impairments in hippocampal function and consequently memory capacities remain to be explored.

## METHODS

### Ethical statement

All animal procedures were performed in accordance with the regulations of the University of Michigan animal care committee.

### Subjects

A total of 20 adult male Sprague Dawley rats (400 - 500 g) were used in this study. Subjects were housed individually in transparent acrylic cages and maintained under controlled temperature (22 ± 1 °C), *ad libitum* food and water, and a 12-h light cycle/12-h dark cycle (lights on at 9:00 am). All subjects were weighed and handled a week prior to any experimental manipulation. For the hippocampal SWR recordings, a tetrodes microdrive array or stainless steel electrodes were used^1^. Briefly, subjects were anesthetized with ketamine (70mg/kg) and xylazine (10 mg/Kg) and kept under 0.5-1.5% of isoflurane the entire time of the surgery. A craniotomy was performed over the dorsal CA1 region of the hippocampus and a microdrive with 8-12 independently moveable tetrodes (four 12.5 μm nichrome wires bundle) or stainless-steel electrodes (two 50 μm wires bundle) were secured in place using dental acrylic with supporting anchoring skull screws. All hippocampal electrodes were dipped in DiI (Sigma Aldrich, St. Louis) prior to surgery for histological analysis. For the electroencephalogram (EEG) recordings, a stainless-steel screw making contact with the dura was placed over the right primary somatosensory cortex (- 1.0 mm AP, 4.0 ML). For the electromyogram (EMG), two stainless steel loops were inserted into the neck muscles. Lastly, a ground screw was placed over the cerebellum (2.0 mm posterior to lambda, 3.5 mm ML). During the course of the surgery, subjects received saline chloride solution (s.c., 5 mL tops) and a carprofen injection (5 mg/Kg, s.c.) was given as an analgesic. All subjects had 7 days of post-surgery recovery. For the subjects implanted with the microdrive, tetrodes were advanced gradually across 11 days post-surgery towards the hippocampus and SWR observation was used as validation for tetrode location. Following recovery, all subjects were acclimated to the middle chamber of the CPP apparatus.

### Electrophysiological recordings

Hippocampal Local Field Potential (LFP), EMG and EEG signals were acquired using a Tucker-Davis Technologies (TDT) acquisition system and Synapse software (TDT). All data were sampled at 6KHz and digitally filtered from 0.5 to 500 Hz and stored for analysis. For all sessions, two video cameras (Allied vision -30fps- and USB Logitech camera -10fps-) were used to monitor the subject’s behavior and awake-sleep state. Video recordings were acquired by RV2 acquisition system (TDT), Synapse software, or open-source VLC software.

#### Online detection of awake and sleep phases

Awake, rest, and sleep states were determined by online recording of EEG and EMG signals with visual supervision (Brown, Basheer, McKenna, Strecker, & McCarley, 2012). Visual criteria used to identify the awake state were the subject’s body movement and spontaneous behavior display (exploring, rearing, grooming, sniffing, and nesting (Maloney, Cape, Gotman, & Jones, 1997). To identify the rest state, the visual criteria used were a reduction of the subject’s body movement or lack of it, together with active head movement, sniffing, occasional grooming, and open eyes. Visual criteria used to identify the sleep state were the subject’s position, immobility for more than 8 seconds, and eyes closed. In addition to these observations, the EMG signal was filtered between 10-100 Hz, and the Root Mean Square (RMS) power averaged within a 5 s sliding window was used to assess the subject’s movement. Sleep state was identified as a decrease in EMG power from 2.5-3.0 std from the mean. To discriminate Rapid Eye Movement (REM) from Non-REM (NREM) sleep phases, the EMG signal had to be below 1.0 std deviation from the mean, indicating the loss of postural muscle tone together with sudden and short power increases produced by muscular twitches. EEG signals were filtered between 0.5-4.5 Hz and 6-10Hz to monitor Delta and Theta activity, respectively. RMS power from both frequency bands was averaged within a 2.5 s sliding window and the delta/theta ratio was calculated. REM and NREM sleep phases were defined by a delta/theta ratio below or near 1.0 and above 2.0, respectively.

#### SWRs real-time detection and sound stimulation

Once criteria for sleep and NREM phase were reached, hippocampal LFP signals from 2 channels with clear SWRs and low spike units were filtered between 150-250 Hz. LFP filtered signals were averaged for 60 s to get the mean basal hippocampal activity. Events from the filtered signals over the mean + 6-9 std in a 10 ms sliding window were considered positive SWRs. These settings were verified to detect events longer than 30 ms. When enabled, each SWR triggered a 50 ms BBN pulse with an intensity of 50 dB and a 5 ms onset and offset ramps (sampling frequency 25 kHz). The SWRs detected from those 2 channels were used for offline analysis of all the electrodes in the hippocampus.

#### Sleep recordings and sound stimulation

Electrophysiological recordings during sleep were made in a chamber (12×12×12 inches, with nesting material as the floor) after spontaneous behavior, defined as the subject’s voluntary behavior (no task performed), or after a hippocampal-dependent associative learning task. During the recordings, the subjects were restricted from water and food, as they may act as distractors. Spontaneous behavior was considered exploring, rearing, grooming, sniffing, and nesting (Maloney et al., 1997). During this session, the subjects were not exposed to any stimuli of any kind. The recording session lasted until it reached 2h of sleep (3-5h total recording time). Sleep was determined by video supervision criteria and electrophysiological signals (as described above). Once the NREM phase was reached, SWRs detection was enabled. The first hour of sleep was recorded without any sound stimulation (No Stimulation -NS-phase). In the second hour, every SWR detection triggered a BBN pulse. Depending on the experimental condition, sound stimuli were not delivered (NS condition), paired to the SWR detection (<5 ms delay, referred to as On-SWR condition), or delayed 2 seconds after the SWR detection (Off-SWR condition).

To test the above following hippocampal-dependent associative learning tasks, subjects were trained in the Conditioned Place Preference (CPP) task and kept in the middle chamber of the apparatus (**Figure 2A**) or in the Contextual Fear Conditioning (CFC) task (**Supplementary Figure 2**) and moved to a different chamber, prior to sleep recordings. The recording session started as soon as the subject slept and once the NREM phase started, the SWR detection was enabled for 2h of sleep (3-4h total recording time). For these recordings, the sound stimulation, either On-SWR or Off-SWR lasted the full 2h of the sleep session.

### Associative learning tasks

#### Conditioned Place Preference (CPP)

The CPP apparatus had three chambers (12×12×12 inches each) divided by two sliding doors. The middle chamber walls were all black and had nesting material as the floor. The chambers on the sides had colored shape patterns posted on the walls and access to a reward well. Each reward well had attached two IR break beam sensors connected to an Arduino Uno REV3 board. The Arduino board was connected to an infusion pump (KDS200 series, KdScientific). On top of the apparatus, one USB Logitech camera (30 fps) was set. Both video recordings and Arduino-automated reward delivery were connected to the TDT RZ2 acquisition system controlled by Synapse software (TDT). When the TDT system was set to enable Arduino’s sensors, every time the subject licked the reward well a TTL sent by Arduino triggered the infusion pump to deliver 0.1 mL of sucrose 20% as a reward at 20mL/min.

CPP behavior paradigm was modified from Trouche et al., 2019 (Trouche et al., 2019). Before training, all the subjects were deprived of water for 24 hours. For training, the subjects were placed in the middle chamber for 1 min with no access to any other chamber. Afterward, the doors accessing the side chambers were open. The subject was allowed to explore the three chambers freely for 15 min (Pre-Test period). During this period, the amount of time spent in each chamber was measured and the innate preference for one of the sided chambers was determined to use the opposite chamber (less time spent in) as the conditioned chamber. After the Pre-Test period, all doors were closed, and the subject was left in the middle chamber for 1 min. To start a conditioning trial, the door to the designated conditioned chamber was opened. After the subject had crossed to the chamber (a full cross was considered as 4 paws inside the chamber), the door was closed. The subject was left in the chamber with access to a solution of 20% sucrose in the reward well. After 5 min, the door was opened to let the subject out of the chamber. Once out, the door was closed again. The subject remained in the middle chamber for 1 min. Next, the opposite door was opened (non-conditioned chamber). The subject remained in the non-conditioned chamber for 5 min with access to plain water in the reward well. Each subject had 2 trials in each chamber in a pseudorandom order. At the end of the trials, the subject was left in the middle chamber to sleep for a period of 3-4h. To test memory retention, two time points were evaluated, a short period (3 h), and a longer period (24 h) after the training. For these tests, subjects were placed in the middle chamber for 1 min. Next, both side doors were opened, and the subject was allowed to move freely through the three chambers for 5 min (Retention period).

#### Contextual Fear Conditioning (CFC)

The CFC apparatus consisted of a transparent acrylic chamber (12×12×12 inches). The chamber had a lateral door and a grid floor of stainless steel rods (Med Associates Inc.). All walls were covered with paper patterns as visual cues. The floor rods could be electrified using a square-pulse stimulator (ENV-414S, Med Associates). On top of the apparatus, a USB Logitech camera (30 fps) was set. Foot shock delivery and video recording were controlled and recorded by Synapse software (TDT). After the CFC conditioning, the subject was removed from the chamber and placed to another transparent acrylic chamber (12×12×12 inches) with white foam board on the walls and nesting material on the floor (“sleep” chamber, Med Associates Inc.).

For the training, the subject was placed inside the conditioning chamber. The subject was allowed to explore the chamber for 5 minutes (pre-Conditioning period). Right after, 6 pulses of 0.45 mA at 0.1 Hz were delivered through the grid floor for one minute (aversive conditioning). Once the foot shock finished, the subject remained in the chamber for another minute (post-conditioning phase). No electrophysiological signals were recording during the CFC conditioning. Afterwards, the subject was removed and placed in the “sleep” chamber to sleep for a period of 3-4h. Memory retention was evaluated 3 h and 24 h after the training. The retention tests consisted of placing the subject in the conditioning chamber for 5 minutes (retention period) with no additional aversive stimuli.

### Histology

To verify the electrodes’ track, all subjects were euthanized under isoflurane anesthesia (5%) and perfused transcardially with saline followed by 4% paraformaldehyde (PFA). Brains were removed and fixed in 4% PFA (72h), followed by cryoprotection in 30% sucrose (72h).

Coronal sections (30-50 μm) were obtained using a cryostat (Leica CM1950) and kept in PBS 4°C before mounting. Each section was mounted on a glass slide and covered with Fluoroshield-DAPI (Abcam, USA). Sections were examined for cell nuclei (DAPI 470 nm) and electrodes track (DiI, 550 nm) using a fluorescence microscope (Zeiss AX10 Imager.M2) and the ZenPro software (Zen (blue edition) version 3.97).

### Data analysis and statistics

To evaluate memory retention in the CPP task, a CPP score was calculated for each pre-Test and retention period (3 h and 24 h post-training). The score was calculated by measuring the amount of time spent in the conditioned chamber minus the amount of time spent in the non-conditioned chamber, divided by the total amount of time spent in both chambers. To get a sense of the absolute change in the subject’s context preference (net preference shift score) and therefore memory robustness, the CPP scores were normalized by subtracting the pre-Test score from the score of the retention period and then dividing by 1 minus the pre-Test score. CPP scores comparison between task’s tests (pre-Test, at 3 h and at 24 h) within a group, were analyzed with the non-parametric repeated-measures Friedman test and the Wilcoxon’s signed-rank test as a post hoc. Net preference shift scores comparison to zero (zero considered as no shift in the contextual preference and therefore, no memory) was analyzed by one-sample Wilcoxon test. Net preference shift scores comparison among groups was analyzed using Kruskal-Wallis and the Mann-Whitney’s U tests. To evaluate aversive memory retention in the CFC task, freezing behavior was calculated during the pre-conditioning period and compared to both retention periods at 3 h and 24 h post-training. Freezing behavior was defined as continuous immobility for more than 5 s, showing no other behavior such as grooming or exploring behavior (actively sniffing in a particular zone). The proportion of the amount of time that the subject spent immobile (% freezing) was calculated by dividing the amount of the time of immobility by the total duration of the test (5 min). Comparison of the percentage of freezing behavior between pre-conditioning, and retention periods (3 h and 24 h tests), was analyzed with the non-parametric repeated-measures Friedman test and the Wilcoxon’s signed-rank test as a post hoc. Comparison of the percentage of freezing behavior between experimental groups was analyzed with Kruskal-Wallis and the Mann-Whitney’s U tests.

#### Awake-sleep states classification and quantification

Electrophysiological recordings were analyzed offline. For awake-sleep states analysis each recording video was visually inspected to choose periods of 900 s where the subject was clearly awake (that included quiet wakefulness states) and sleeping. For all 900 s bouts per state, both EMG and EEG signals were filtered from 10-100 Hz and 0.5-30 Hz, respectively. First, to analyze subject’s movement per state, EMG power was calculated using the RMS in a 10 s window. Based on the EMG power comparison between states a threshold of the EMG power + 1.5 std deviation was used for awake-sleep states classification for the following analysis of the recording session. Proportion of the total time per state per session was calculated by dividing the amount of time per state by the total time of the recording session. Proportion of EMG power during sleep was normalized to the EMG power during awake. To classify NREM and REM sleep phases, the EEG signal was filtered between 0.5-4.5Hz corresponding to the Delta frequency band and between 6-10 Hz corresponding to the Theta frequency band. Delta and Theta power were calculated using the RMS in a 2.5 s window. Periods with Delta/Theta ratios below 1.0 were classified as REM periods, while periods with Delta/Theta ratios higher than 2.0 were classified as NREM periods. To get the proportion of the NREM and REM periods, the amount of time in NREM and REM periods was normalized to the total amount of sleep per recording session.

#### Real-time detected SWRs validation and analysis

Putative SWRs detected during the recording acquisition were reevaluated offline to verify their identity and remove any potential false positives. The criteria of SWRs identification were to have a ripple power greater than the mean ripple power + 3 std deviation and a duration greater than 30 ms. To do this validation, the hippocampal LFP signal from NREM periods of sleep was band-pass filtered between 150 and 250 Hz. Next, the mean and the threshold of the ripple power was calculated. Ripple power from each event detected was evaluated in windows of 100 ms before and after the event-detected time. The events having a ripple power over the threshold were selected. Next, high ripple power events duration was calculated using the ripple power autocorrelation. Ripple duration was estimated from autocorrelations peak to periods where the autocorrelation peaks remained above the autocorrelation’s mean ± 2.0 std deviation (Ellender, Nissen, Colgin, Mann, & Paulsen, 2010). Events with a duration shorter than 30 ms were discarded. Next, spectrograms (0 - 400 Hz) were calculated for each 200 ms window from verified SWRs (-100 ms, +100 ms, from SWR detection). Then, spectrograms were normalized by the mean power spectrogram of the whole recording from that session. From the normalized spectrograms, individual SWR peak frequency and power in time domain were quantified. Individual SWRs normalized ripple power was quantified from 150-250 Hz and from 0 - 100 ms from SWRs detection. Finally, spectrograms from individual SWRs were averaged to get the mean spectrogram per recording. Spectrograms were averaged to get a mean spectrogram per experimental condition. SWRs rate was averaged per subject while ripple’s power, duration and peak frequency were averaged grouping SWRs per experimental condition.

#### periSWRs EMG analysis

To further verify the effect of the sound stimulation over the awake-sleep states, averaged EMG power and Delta power of 5s windows were compared before and after sound stimulation times. Comparison between EMG power or Delta power from Pre-sound stimulus windows to Post-sound stimulus windows were done by getting the logarithm of the ratio from EMG power or Delta power Pre/Post and compared to zero using one sample Wilcoxon’s signed-rank test.

EEG, EMG, and SWRs data (rate, power, duration, and peak frequency) were analyzed with non-parametric statistics. To compare the effect of the sound stimulation within a group, data were analyzed with two-tailed Wilcoxon’s signed-rank test (WSR). To compare the effect of sound stimulation between two groups Mann-Whitney’s U (MW-U) test was used. To compare the effect of sound stimulation among groups, Kruskal-Wallis (KW) analysis was used followed by the post hoc Tukey-Kramer (T-K) test. SWRs power over time waves were analyzed with a two-way ANOVA.

Data and statistical analysis were performed with MATLAB custom scripts and standard functions (R2019a). Figures were made using Adobe Illustrator software (CS6 v16.0.0). Figures schemes were created using the BioRender.com platform.

## AUTHORS CONTRIBUTIONS

G.R. and K.S-P. conceptualized and designed the study, K.S-P. conducted experiments, K.S-P. analyzed the data, K.S-P. and G.R. wrote the manuscript

## ACKNOWLEDGEMENTS

This work was supported by a National Institute of Health grant R01NS129874 (G.R.), a National Institute of Health grant R01MH063649 (G.R. Co-I) and an Alzheimer’s Association Research Grant 21-850571 (G.R.)

## REFERENCES

Antony, J. W., Gobel, E. W., O’Hare, J. K., Reber, P. J., & Paller, K. A. (2012). Cued memory reactivation during sleep influences skill learning. Nature Neuroscience, 15(8), 1114-+. doi:10.1038/nn.3152

Aronov, D., Nevers, R., & Tank, D. W. (2017). Mapping of a non-spatial dimension by the hippocampal-entorhinal circuit. Nature, 543(7647), 719–722. doi:10.1038/nature21692

Bar, E., Marmelshtein, A., Arzi, A., Perl, O., Livne, E., Hizmi, E., . . . Nir, Y. (2020). Local Targeted Memory Reactivation in Human Sleep. Current Biology, 30(8), 1435–1446 e1435. doi:10.1016/j.cub.2020.01.091

Batterink, L. J., Creery, J. D., & Paller, K. A. (2016). Phase of Spontaneous Slow Oscillations during Sleep Influences Memory-Related Processing of Auditory Cues. The Journal of neuroscience : the official journal of the Society for Neuroscience, 36(4), 1401–1409. doi:10.1523/jneurosci.3175-15.2016

Bendor, D., & Wilson, M. A. (2012). Biasing the content of hippocampal replay during sleep. Nature Neuroscience, 15(10), 1439–1444. doi:10.1038/nn.3203

Billig, A. J., Lad, M., Sedley, W., & Griffiths, T. D. (2022). The hearing hippocampus. Prog Neurobiol, 218, 102326. doi:10.1016/j.pneurobio.2022.102326

Borg, E. (1982). Auditory thresholds in rats of different age and strain. A behavioral and electrophysiological study. Hear Res, 8(2), 101–115. doi:10.1016/0378-5955(82)90069-7

Brink, M., Omlin, S., Muller, C., Pieren, R., & Basner, M. (2011). An event-related analysis of awakening reactions due to nocturnal church bell noise. Science of the Total Environment, 409(24), 5210–5220. doi:10.1016/j.scitotenv.2011.09.020

Brodt, S., Inostroza, M., Niethard, N., & Born, J. (2023). Sleep-A brain-state serving systems memory consolidation. Neuron, 111(7), 1050–1075. doi:10.1016/j.neuron.2023.03.005

Brown, R. E., Basheer, R., McKenna, J. T., Strecker, R. E., & McCarley, R. W. (2012). Control of Sleep and Wakefulness. Physiological Reviews, 92(3), 1087–1187. doi:10.1152/physrev.00032.2011

Budinger, E., & Scheich, H. (2009). Anatomical connections suitable for the direct processing of neuronal information of different modalities via the rodent primary auditory cortex. Hear Res, 258(1-2), 16–27. doi:10.1016/j.heares.2009.04.021

Buzsaki, G. (1989). Two-stage model of memory trace formation: a role for “noisy” brain states. Neuroscience, 31(3), 551–570. doi:10.1016/0306-4522(89)90423-5

Buzsaki, G. (2015). Hippocampal sharp wave-ripple: A cognitive biomarker for episodic memory and planning. Hippocampus, 25(10), 1073–1188. doi:10.1002/hipo.22488

Cellini, N., & Capuozzo, A. (2018). Shaping memory consolidation via targeted memory reactivation during sleep. Annals of the New York Academy of Sciences, 1426(1), 52–71. doi:10.1111/nyas.13855

Chambers, A. R., Berge, C. N., & Vervaeke, K. (2022). Cell-type-specific silence in thalamocortical circuits precedes hippocampal sharp-wave ripples. Cell Reports, 40(4). doi:ARTN 111132 10.1016/j.celrep.2022.111132

Chepesiuk, R. (2005). Decibel hell: The effects of living in a noisy world. Environmental Health Perspectives, 113(1), A34–A41. doi:DOI 10.1289/ehp.113-a34

Creery, J. D., Brang, D. J., Arndt, J. D., Bassard, A., Towle, V. L., Tao, J. X., . . . Paller, K. A. (2022). Electrophysiological markers of memory consolidation in the human brain when memories are reactivated during sleep. Proc Natl Acad Sci U S A, 119(44), e2123430119. doi:10.1073/pnas.2123430119

Creery, J. D., Oudiette, D., Antony, J. W., & Paller, K. A. (2015). Targeted Memory Reactivation during Sleep Depends on Prior Learning. Sleep, 38(5), 755–763. doi:10.5665/sleep.4670

Diba, K., & Buzsaki, G. (2007). Forward and reverse hippocampal place-cell sequences during ripples. Nature Neuroscience, 10(10), 1241–1242. doi:10.1038/nn1961

Diekelmann, S., & Born, J. (2010). The memory function of sleep. Nat Rev Neurosci, 11(2), 114–126. doi:10.1038/nrn2762

Edeline, J. M., Dutrieux, G., Manunta, Y., & Hennevin, E. (2001). Diversity of receptive field changes in auditory cortex during natural sleep. European Journal of Neuroscience, 14(11), 1865–1880. doi:10.1046/j.0953-816x.2001.01821.x

Ego-Stengel, V., & Wilson, M. A. (2010). Disruption of ripple-associated hippocampal activity during rest impairs spatial learning in the rat. Hippocampus, 20(1), 1–10. doi:10.1002/hipo.20707

Ellender, T. J., Nissen, W., Colgin, L. L., Mann, E. O., & Paulsen, O. (2010). Priming of Hippocampal Population Bursts by Individual Perisomatic-Targeting Interneurons. Journal of Neuroscience, 30(17), 5979–5991. doi:10.1523/jneurosci.3962-09.2010

Eschenko, O., Ramadan, W., Molle, M., Born, J., & Sara, S. J. (2008). Sustained increase in hippocampal sharp-wave ripple activity during slow-wave sleep after learning. Learn Mem, 15(4), 222–228. doi:10.1101/lm.726008

Europe, W. H. O. R. O. f., World Health Organization, E., Hurtley, C., & World Health Organization. Regional Office for, E. (2009). Night noise guidelines for Europe. Copenhagen: World Health Organization Europe.

Fernandez-Ruiz, A., Oliva, A., Fermino de Oliveira, E., Rocha-Almeida, F., Tingley, D., & Buzsaki, G. (2019). Long-duration hippocampal sharp wave ripples improve memory. Science, 364(6445), 1082–1086. doi:10.1126/science.aax0758

Fiedler, P. E. K., & Zannin, P. H. T. (2015). Evaluation of noise pollution in urban traffic hubs-Noise maps and measurements. Environmental Impact Assessment Review, 51, 1–9. doi:10.1016/j.eiar.2014.09.014

Foster, D. J. (2017). Replay Comes of Age. Annu Rev Neurosci, 40, 581–602. doi:10.1146/annurev-neuro-072116-031538

Frankland, P. W., & Bontempi, B. (2005). The organization of recent and remote memories. Nat Rev Neurosci, 6(2), 119–130. doi:10.1038/nrn1607

Fritschi, L., & World health Organization. Regional Office for, E. (2011). Burden of disease from environmental noise : quantification of healthy life years lost in Europe. Denmark: World Health Organization, Regional Office for Europe.

Furtak, S. C., Wei, S. M., Agster, K. L., & Burwell, R. D. (2007). Functional neuroanatomy of the parahippocampal region in the rat: the perirhinal and postrhinal cortices. Hippocampus, 17(9), 709–722. doi:10.1002/hipo.20314

Geva-Sagiv, M., Mankin, E. A., Eliashiv, D., Epstein, S., Cherry, N., Kalender, G., . . . Fried, I. (2023). Augmenting hippocampal-prefrontal neuronal synchrony during sleep enhances memory consolidation in humans. Nature Neuroscience, 26(6), 1100–1110. doi:10.1038/s41593-023-01324-5

Geva-Sagiv, M., & Nir, Y. (2019). Local Sleep Oscillations: Implications for Memory Consolidation. Front Neurosci, 13, 813. doi:10.3389/fnins.2019.00813

Girardeau, G., Benchenane, K., Wiener, S. I., Buzsaki, G., & Zugaro, M. B. (2009). Selective suppression of hippocampal ripples impairs spatial memory. Nature Neuroscience, 12(10), 1222–1223. doi:10.1038/nn.2384

Girardeau, G., Cei, A., & Zugaro, M. (2014). Learning-Induced Plasticity Regulates Hippocampal Sharp Wave-Ripple Drive. Journal of Neuroscience, 34(15), 5176–5183. doi:10.1523/Jneurosci.4288-13.2014

Graves, L. A., Heller, E. A., Pack, A. I., & Abel, T. (2003). Sleep Deprivation Selectively Impairs Memory Consolidation for Contextual Fear Conditioning. Learning \& Memory, 10(3), 168–176. doi:10.1101/lm.48803

Gridchyn, I., Schoenenberger, P., O’Neill, J., & Csicsvari, J. (2020). Assembly-Specific Disruption of Hippocampal Replay Leads to Selective Memory Deficit. Neuron. doi:10.1016/j.neuron.2020.01.021

Hagewoud, R., Whitcomb, S. N., Heeringa, A. N., Havekes, R., Koolhaas, J. M., & Meerlo, P. (2010). A Time for Learning and a Time for Sleep: The Effect of Sleep Deprivation on Contextual Fear Conditioning at Different Times of the Day. Sleep, 33(10), 1315–1322. doi:10.1093/sleep/33.10.1315

Hahad, O., Jimenez, M. T. B., Kuntic, M., Frenis, K., Steven, S., Daiber, A., & Münzel, T. (2022). Cerebral consequences of environmental noise exposure. Environment International, 165, 107306. doi:10.1016/j.envint.2022.107306

Hasselmo, M. E. (2003). Theta theory: Requirements for encoding events and task rules explain theta phase relationships in hippocampus and neocortex. Proceedings of the International Joint Conference on Neural Networks, 2003, 2, 1470–1475. doi:10.1109/ijcnn.2003.1223914

Hasselmo, M. E., & Stern, C. E. (2013). Theta rhythm and the encoding and retrieval of space and time. NeuroImage, 85. doi:10.1016/j.neuroimage.2013.06.022

Hayat, H., Marmelshtein, A., Krom, A. J., Sela, Y., Tankus, A., Strauss, I., Nir, Y. (2022). Reduced neural feedback signaling despite robust neuron and gamma auditory responses during human sleep. Nat Neurosci, 25(7), 935–943. doi:10.1038/s41593-022-01107-4

Insausti, R., Herrero, M. T., & Witter, M. P. (1997). Entorhinal cortex of the rat: cytoarchitectonic subdivisions and the origin and distribution of cortical efferents. Hippocampus, 7(2), 146–183. doi:10.1002/(SICI)1098-1063(1997)7:2<146::AID-HIPO4>3.0.CO;2-L

Ip, Z., Rabiller, G., He, J. W., Chavan, S., Nishijima, Y., Akamatsu, Y., Yazdan-Shahmorad, A. (2021). Local field potentials identify features of cortico-hippocampal communication impacted by stroke and environmental enrichment therapy. Journal of Neural Engineering, 18(4). doi:ARTN 0460a1 10.1088/1741-2552/ac0a54

Issa, E. B., & Wang, X. (2008). Sensory responses during sleep in primate primary and secondary auditory cortex. J Neurosci, 28(53), 14467–14480. doi:10.1523/JNEUROSCI.3086-08.2008

Itskov, P. M., Vinnik, E., Honey, C., Schnupp, J., & Diamond, M. E. (2012). Sound sensitivity of neurons in rat hippocampus during performance of a sound-guided task. J Neurophysiol, 107(7), 1822–1834. doi:10.1152/jn.00404.2011

Izquierdo, I., Barros, D. M., Mello e Souza, T., de Souza, M. M., Izquierdo, L. A., & Medina, J. H. (1998). Mechanisms for memory types differ. Nature, 393(6686), 635–636. doi:10.1038/31371

Izquierdo, L. A., Barros, D. M., Ardenghi, P. G., Pereira, P., Rodrigues, C., Choi, H., Izquierdo, I. (2000). Different hippocampal molecular requirements for short- and long-term retrieval of one-trial avoidance learning. Behavioural Brain Research, 111(1-2), 93–98. doi:10.1016/s0166-4328(00)00137-6

Izquierdo, L. A., Vianna, M., Barros, D. M., Souza, T. M. e., Ardenghi, P., Anna, M. K. S., Izquierdo, I. (2000). Short- and Long-Term Memory Are Differentially Affected by Metabolic Inhibitors Given into Hippocampus and Entorhinal Cortex. Neurobiology of Learning and Memory, 73(2), 141–149. doi:10.1006/nlme.1999.3925

Jadhav, S. P., Kemere, C., German, P. W., & Frank, L. M. (2012). Awake hippocampal sharp-wave ripples support spatial memory. Science, 336(6087), 1454–1458. doi:10.1126/science.1217230

Jadhav, S. P., Rothschild, G., Roumis, D. K., & Frank, L. M. (2016). Coordinated Excitation and Inhibition of Prefrontal Ensembles during Awake Hippocampal Sharp-Wave Ripple Events. Neuron, 90(1), 113–127. doi:10.1016/j.neuron.2016.02.010

Jarosinska, D., Heroux, M. E., Wilkhu, P., Creswick, J., Verbeek, J., Wothge, J., & Paunovic, E. (2018). Development of the WHO Environmental Noise Guidelines for the European Region: An Introduction. Int J Environ Res Public Health, 15(4). doi:10.3390/ijerph15040813

Jarzebowski, P., Tang, C. S., Paulsen, O., & Hay, Y. A. (2021). Impaired spatial learning and suppression of sharp wave ripples by cholinergic activation at the goal location. Elife, 10. doi:ARTN e65998 10.7554/eLife.65998

Ji, D., & Wilson, M. A. (2007). Coordinated memory replay in the visual cortex and hippocampus during sleep. Nat Neurosci, 10(1), 100–107. doi:10.1038/nn1825

Kerr, K. M., Agster, K. L., Furtak, S. C., & Burwell, R. D. (2007). Functional neuroanatomy of the parahippocampal region: the lateral and medial entorhinal areas. Hippocampus, 17(9), 697–708. doi:10.1002/hipo.20315

Kraus, K. S., & Canlon, B. (2012). Neuronal connectivity and interactions between the auditory and limbic systems. Effects of noise and tinnitus. Hearing Research, 288(1-2), 34–46. doi:10.1016/j.heares.2012.02.009

Kronenberg, P. v., Milinski, L., Kruschke, Z., & Hoz, L. d. (2022). Sound disrupts sleep-associated brain oscillations in rodents in a meaning-dependent manner. Scientific Reports, 12(1), 6051. doi:10.1038/s41598-022-09457-6

Lee, J. L., Everitt, B. J., & Thomas, K. L. (2004). Independent cellular processes for hippocampal memory consolidation and reconsolidation. Science, 304(5672), 839–843. doi:10.1126/science.1095760

Leminen, M., Virkkala, J., Saure, E., Paajanen, T., Zee, P., Santostasi, G., . . . Paunio, T. (2017). Enhanced Memory Consolidation Via Automatic Sound Stimulation during Non-REM Sleep. Sleep, 40(3), zsx003. doi:10.1093/sleep/zsx003

Lewis, P. A., & Bendor, D. (2019). How Targeted Memory Reactivation Promotes the Selective Strengthening of Memories in Sleep. Current Biology, 29(18), R906–R912. doi:10.1016/j.cub.2019.08.019

Maloney, K. J., Cape, E. G., Gotman, J., & Jones, B. E. (1997). High-frequency γ electroencephalogram activity in association with sleep-wake states and spontaneous behaviors in the rat. Neuroscience, 76(2), 541–555. doi:10.1016/s0306-4522(96)00298-9

Mascagni, F., McDonald, A. J., & Coleman, J. R. (1993). Corticoamygdaloid and corticocortical projections of the rat temporal cortex: a Phaseolus vulgaris leucoagglutinin study. Neuroscience, 57(3), 697–715. doi:10.1016/0306-4522(93)90016-9

Memmesheimer, R.-M. (2010). Quantitative prediction of intermittent high-frequency oscillations in neural networks with supralinear dendritic interactions. Proceedings of the National Academy of Sciences, 107(24), 11092–11097. doi:10.1073/pnas.0909615107

Michon, F., Sun, J.-J., Kim, C. Y., Ciliberti, D., & Kloosterman, F. (2019). Post-learning Hippocampal Replay Selectively Reinforces Spatial Memory for Highly Rewarded Locations. Current Biology, 29(9), 1436–1444.e1435. doi:10.1016/j.cub.2019.03.048

Miller, C. L., & Freedman, R. (1993). Medial septal neuron activity in relation to an auditory sensory gating paradigm. Neuroscience, 55(2), 373–380. doi:10.1016/0306-4522(93)90506-b

Moxon, K. A., Gerhardt, G. A., Bickford, P. C., Austin, K., Rose, G. M., Woodward, D. J., & Adler, L. E. (1999). Multiple single units and population responses during inhibitory gating of hippocampal auditory response in freely-moving rats. Brain Res, 825(1-2), 75–85. doi:10.1016/s0006-8993(99)01187-7

Nir, Y., Vyazovskiy, V. V., Cirelli, C., Banks, M. I., & Tononi, G. (2015). Auditory responses and stimulus-specific adaptation in rat auditory cortex are preserved across NREM and REM sleep. Cereb Cortex, 25(5), 1362–1378. doi:10.1093/cercor/bht328

Norimoto, H., Makino, K., Gao, M., Shikano, Y., Okamoto, K., Ishikawa, T., . . . Ikegaya, Y. (2018). Hippocampal ripples down-regulate synapses. Science, 359(6383), 1524–1527. doi:10.1126/science.aao0702

O’Keefe, J., & Nadel, L. (1978). The hippocampus as a cognitive map. Oxford New York: Clarendon Press ; Oxford University Press.

Oudiette, D., & Paller, K. A. (2013). Upgrading the sleeping brain with targeted memory reactivation. Trends in Cognitive Sciences, 17(3), 142–149. doi:10.1016/j.tics.2013.01.006

Paul, K. C., Haan, M., Mayeda, E. R., & Ritz, B. R. (2019). Ambient Air Pollution, Noise, and Late-Life Cognitive Decline and Dementia Risk. Annual Review of Public Health, 40(1), 203–220. doi:10.1146/annurev-publhealth-040218-044058

Pena, J. L., Perez-Perera, L., Bouvier, M., & Velluti, R. A. (1999). Sleep and wakefulness modulation of the neuronal firing in the auditory cortex of the guinea pig. Brain Res, 816(2), 463–470. doi:10.1016/s0006-8993(98)01194-9

Peyrache, A., Khamassi, M., Benchenane, K., Wiener, S. I., & Battaglia, F. P. (2009). Replay of rule-learning related neural patterns in the prefrontal cortex during sleep. Nature Neuroscience, 12(7), 919–926. doi:10.1038/nn.2337

Pignatelli, M., Ryan, T. J., Roy, D. S., Lovett, C., Smith, L. M., Muralidhar, S., & Tonegawa, S. (2019). Engram Cell Excitability State Determines the Efficacy of Memory Retrieval. Neuron, 101(2), 274–284.e275. doi:10.1016/j.neuron.2018.11.029

Pinto, L., Rajan, K., DePasquale, B., Thiberge, S. Y., Tank, D. W., & Brody, C. D. (2019). Task-Dependent Changes in the Large-Scale Dynamics and Necessity of Cortical Regions. Neuron, 104(4), 810–824.e819. doi:10.1016/j.neuron.2019.08.025

Poo, M.-M. M., Pignatelli, M., Ryan, T. J. J., Tonegawa, S., Bonhoeffer, T., Martin, K. C., . . . Stevens, C. (2016). What is memory? The present state of the engram. BMC biology, 14(1), 40. doi:10.1186/s12915-016-0261-6

Purple, R. J., Sakurai, T., & Sakaguchi, M. (2017). Auditory conditioned stimulus presentation during NREM sleep impairs fear memory in mice. Scientific Reports, 7(1), 46247. doi:10.1038/srep46247

Redondo, R. L., Kim, J., Arons, A. L., Ramirez, S., Liu, X., & Tonegawa, S. (2014). Bidirectional switch of the valence associated with a hippocampal contextual memory engram. Nature, 513(7518), 426–430. doi:10.1038/nature13725

Rothschild, G., Eban, E., & Frank, L. M. (2017). A cortical-hippocampal-cortical loop of information processing during memory consolidation. Nature Neuroscience, 20(2), 251–259. doi:10.1038/nn.4457

Roy, D. S., Kitamura, T., Okuyama, T., Ogawa, S. K., Sun, C., Obata, Y., . . . Tonegawa, S. (2017). Distinct Neural Circuits for the Formation and Retrieval of Episodic Memories. Cell, 170(5), 1000–1012.e1019. doi:10.1016/j.cell.2017.07.013

Rudoy, J. D., Voss, J. L., Westerberg, C. E., & Paller, K. A. (2009). Strengthening Individual Memories by Reactivating Them During Sleep. Science, 326(5956), 1079–1079. doi:10.1126/science.1179013

Rudy, J. W., & Sutherland, R. J. (2008). Is it systems or cellular consolidation? Time will tell. An alternative interpretation of the Morris group’s recent science paper. Neurobiol Learn Mem, 89(4), 366–369. doi:10.1016/j.nlm.2007.07.017

Sadowski, Josef H. L. P., Jones, Matthew W., & Mellor, Jack R. (2016). Sharp-Wave Ripples Orchestrate the Induction of Synaptic Plasticity during Reactivation of Place Cell Firing Patterns in the Hippocampus. Cell Reports, 14(8), 1916–1929. doi:10.1016/j.celrep.2016.01.061

Sakurai, Y. (2002). Coding of auditory temporal and pitch information by hippocampal individual cells and cell assemblies in the rat. Neuroscience, 115(4), 1153–1163. doi:10.1016/s0306-4522(02)00509-2

Schafe, G. E., Nadel, N. V., Sullivan, G. M., Harris, A., & LeDoux, J. E. (1999a). Memory consolidation for contextual and auditory fear conditioning is dependent on protein synthesis, PKA, and MAP kinase. Learn Mem, 6(2), 97–110. Retrieved from https://www.ncbi.nlm.nih.gov/pubmed/10327235

Schafe, G. E., Nadel, N. V., Sullivan, G. M., Harris, A., & LeDoux, J. E. (1999b). Memory consolidation for contextual and auditory fear conditioning is dependent on protein synthesis, PKA, and MAP kinase. Learning & Memory, 6(2), 97–110. Retrieved from <Go to ISI>://WOS:000079905700003

Schechtman, E., Antony, J. W., Lampe, A., Wilson, B. J., Norman, K. A., & Paller, K. A. (2021). Multiple memories can be simultaneously reactivated during sleep as effectively as a single memory. Commun Biol, 4(1), 25. doi:10.1038/s42003-020-01512-0

Schlüter, H., Hackländer, R. P., & Bermeitinger, C. (2019). Emotional oddball: A review on memory effects. Psychonomic Bulletin \& Review, 26(5), 1472–1502. doi:10.3758/s13423-019-01658-x

Schomburg, Erik W., Fernández-Ruiz, A., Mizuseki, K., Berényi, A., Anastassiou, Costas A., Koch, C., & Buzsáki, G. (2014). Theta Phase Segregation of Input-Specific Gamma Patterns in Entorhinal-Hippocampal Networks. Neuron, 84(2), 470–485. doi:10.1016/j.neuron.2014.08.051

Scoville, W. B., & Milner, B. (1957). Loss of recent memory after bilateral hippocampal lesions. J Neurol Neurosurg Psychiatry, 20(1), 11–21. doi:10.1136/jnnp.20.1.11

Sela, Y., Krom, A. J., Bergman, L., Regev, N., & Nir, Y. (2020). Sleep Differentially Affects Early and Late Neuronal Responses to Sounds in Auditory and Perirhinal Cortices. J Neurosci, 40(14), 2895–2905. doi:10.1523/JNEUROSCI.1186-19.2020

Skaggs, W. E., & McNaughton, B. L. (1996). Replay of neuronal firing sequences in rat hippocampus during sleep following spatial experience. Science, 271(5257), 1870–1873. doi:10.1126/science.271.5257.1870

Squire, L. R., Genzel, L., Wixted, J. T., & Morris, R. G. (2015). Memory consolidation. Cold Spring Harb Perspect Biol, 7(8), a021766. doi:10.1101/cshperspect.a021766

Steward, O. (1976). Topographic organization of the projections from the entorhinal area to the hippocampal formation of the rat. J Comp Neurol, 167(3), 285–314. doi:10.1002/cne.901670303

Trouche, S., Koren, V., Doig, N. M., Ellender, T. J., El-Gaby, M., Lopes-dos-Santos, V., . . . Dupret, D. (2019). A Hippocampus-Accumbens Tripartite Neuronal Motif Guides Appetitive Memory in Space. Cell, 176(6), 1393–1406.e1316. doi:10.1016/j.cell.2018.12.037

Tse, D., Langston, R. F., Kakeyama, M., Bethus, I., Spooner, P. A., Wood, E. R., . . . Morris, R. G. (2007). Schemas and memory consolidation. Science, 316(5821), 76–82. doi:10.1126/science.1135935

Tzivian, L., Dlugaj, M., Winkler, A., Weinmayr, G., Hennig, F., Fuks, K. B., . . . Heinz Nixdorf Recall study Investigative, G. (2016). Long-Term Air Pollution and Traffic Noise Exposures and Mild Cognitive Impairment in Older Adults: A Cross-Sectional Analysis of the Heinz Nixdorf Recall Study. Environ Health Perspect, 124(9), 1361–1368. doi:10.1289/ehp.1509824

Tzivian, L., Jokisch, M., Winkler, A., Weimar, C., Hennig, F., Sugiri, D., . . . Grp, H. N. R. S. (2017). Associations of long-term exposure to air pollution and road traffic noise with cognitive function-An analysis of effect measure modification. Environment International, 103, 30–38. doi:10.1016/j.envint.2017.03.018

van de Ven, G. M., Trouche, S., McNamara, C. G., Allen, K., & Dupret, D. (2016). Hippocampal Offline Reactivation Consolidates Recently Formed Cell Assembly Patterns during Sharp Wave-Ripples. Neuron, 92(5), 968–974. doi:10.1016/j.neuron.2016.10.020

Vandecasteele, M., Varga, V., Berenyi, A., Papp, E., Bartho, P., Venance, L., . . . Buzsaki, G. (2014). Optogenetic activation of septal cholinergic neurons suppresses sharp wave ripples and enhances theta oscillations in the hippocampus. Proceedings of the National Academy of Sciences of the United States of America, 111(37), 13535–13540. doi:10.1073/pnas.1411233111

Varela, C., Kumar, S., Yang, J. Y., & Wilson, M. A. (2014). Anatomical substrates for direct interactions between hippocampus, medial prefrontal cortex, and the thalamic nucleus reuniens. Brain Structure & Function, 219(3), 911–929. doi:10.1007/s00429-013-0543-5

Varela, C., & Wilson, M. A. (2020a). mPFC Spindle Cycles Organize Sparse Thalamic Activation and Recently Active CA1 cells During non-REM Sleep. bioRxiv, 653436. doi:10.1101/653436

Varela, C., & Wilson, M. A. (2020b). mPFC spindle cycles organize sparse thalamic activation and recently active CA1 cells during non-REM sleep. Elife, 9. doi:10.7554/eLife.48881

Velluti, R. A. (1997). Interactions between sleep and sensory physiology. Journal of Sleep Research, 6(2), 61–77. doi:DOI 10.1046/j.1365-2869.1997.00031.x

Vianna, M. R. M., Barros, D. M., Silva, T., Choi, H., Madche, C., Rodrigues, C., . . . Izquierdo, I. (2000). Pharmacological demonstration of the differential involvement of protein kinase C isoforms in short- and long-term memory formation and retrieval of one-trial avoidance in rats. Psychopharmacology, 150(1), 77–84. doi:10.1007/s002130000396

Walz, R., Roesler, R., Quevedo, J., Sant’Anna, M. K., Madruga, M., Rodrigues, C., . . . Izquierdo, I. (2000). Time-Dependent Impairment of Inhibitory Avoidance Retention in Rats by Posttraining Infusion of a Mitogen-Activated Protein Kinase Kinase Inhibitor into Cortical and Limbic Structures. Neurobiology of Learning and Memory, 73(1), 11–20. doi:10.1006/nlme.1999.3913

Warden, M. R., & Miller, E. K. (2010). Task-Dependent Changes in Short-Term Memory in the Prefrontal Cortex. The Journal of Neuroscience, 30(47), 15801–15810. doi:10.1523/jneurosci.1569-10.2010

Weuve, J., D’Souza, J., Beck, T., Evans, D. A., Kaufman, J. D., Rajan, K. B., . . . Adar, S. D. (2020). Long-term community noise exposure in relation to dementia, cognition, and cognitive decline in older adults. Alzheimers Dement. doi:10.1002/alz.12191

Wierzynski, C. M., Lubenov, E. V., Gu, M., & Siapas, A. G. (2009). State-dependent spike-timing relationships between hippocampal and prefrontal circuits during sleep. Neuron, 61(4), 587–596. doi:10.1016/j.neuron.2009.01.011

Wilson, M. A., & McNaughton, B. L. (1994). Reactivation of hippocampal ensemble memories during sleep. Science, 265(5172), 676–679. doi:10.1126/science.8036517

Xiao, C., Liu, Y., Xu, J., Gan, X., & Xiao, Z. (2018). Septal and Hippocampal Neurons Contribute to Auditory Relay and Fear Conditioning. Front Cell Neurosci, 12, 102. doi:10.3389/fncel.2018.00102

Ylinen, A., Bragin, A., Nadasdy, Z., Jando, G., Szabo, I., Sik, A., & Buzsaki, G. Ylinen, 1995.pdf. doi:10.1523/jneurosci.15-01-00030.1995

Zhang, G. W., Sun, W. J., Zingg, B., Shen, L., He, J. F., Xiong, Y., . . . Zhang, L. I. (2018). A Non-canonical Reticular-Limbic Central Auditory Pathway via Medial Septum Contributes to Fear Conditioning. Neuron, 97(2), 406-+. doi:10.1016/j.neuron.2017.12.010

